# Multiscale three-dimensional imaging of intact human organs down to the cellular scale using hierarchical phase-contrast tomography

**DOI:** 10.1101/2021.02.03.429481

**Authors:** C. Walsh, P. Tafforeau, Willi L. Wagner, D. J. Jafree, A. Bellier, C. Werlein, M. P. Kühnel, E. Boller, S. Walker-Samuel, J. L. Robertus, D. A. Long, J. Jacob, S. Marussi, E. Brown, N. Holroyd, D. D. Jonigk, M. Ackermann, P. D. Lee

## Abstract

Human organs are complex, three-dimensional and multiscale systems. Spatially mapping the human body down through its hierarchy, from entire organs to their individual functional units and specialised cells, is a major obstacle to fully understanding health and disease. To meet this challenge, we developed hierarchical phase-contrast tomography (HiP-CT), an X-ray phase propagation technique utilising the European Synchrotron Radiation Facility’s Extremely Brilliant Source: the world’s first high-energy 4^th^ generation X-ray source. HiP-CT enabled three-dimensional and non-destructive imaging at near-micron resolution in soft tissues at one hundred thousand times the voxel size whilst maintaining the organ’s structure. We applied HiP-CT to image five intact human parenchymal organs: brain, lung, heart, kidney and spleen. These were hierarchically assessed with HiP-CT, providing a structural overview of the whole organ alongside detail of the organ’s individual functional units and cells. The potential applications of HiP-CT were demonstrated through quantification and morphometry of glomeruli in an intact human kidney, and identification of regional changes to the architecture of the air-tissue interface and alveolar morphology in the lung of a deceased COVID-19 patient. Overall, we show that HiP-CT is a powerful tool which can provide a comprehensive picture of structural information for whole intact human organs, encompassing precise details on functional units and their constituent cells to better understand human health and disease.

## INTRODUCTION

Biological tissues are inherently complex, three-dimensional (3D) and hierarchically arranged in an ordered series of length scales, ranging from individual specialised cells to larger tissues and then complete organs. Spatial relationships, 3D morphology and interaction within and across these length scales collectively provide a basis for biological function. Thus, mapping the spatial organisation and morphology of individual cells up to the scale of intact organs is fundamental towards understanding system-level behaviours in health or disease. However, the predominant method used to image multi-scale tissues in the life sciences involves physical sub-sampling into smaller sub-sections of tissue, before high-resolution imaging ^1^. A sub-sampling is not ideal for multi-scale imaging, due to the challenges of co-registering data across different modalities and an inability to obtain representative data across all length scales within human organs ^1,2^. Novel 3D imaging techniques are therefore required to bridge length scales from cellular level spatial relationships to the architectural organisation of intact organs.

In recent years there has been progress towards achieving 3D imaging of intact organs at multiple length scales. One approach is optical clearing which homogenises refractive index in biological tissues, allowing high-resolution 3D imaging modalities such as light sheet fluorescence microscopy or optical projection tomography. This has allowed the visualisation of cellular structures at the scale of whole organs or organisms in contexts such as human embryonic development ^3,4^, prediction of organ-wide or organism-wide distribution of metastasis ^5^ and the uptake of drugs ^6^ in mouse models of cancer. Optical clearing of whole adult human organs has recently been achieved ^7^, however, it requires long durations (~3 months) to obtain cleared tissues and causes changes in tissue morphology during preparation. Conversely, high resolution magnetic resonance imaging (MRI) requires minimal tissue preparation and does not cause tissue distortion; achieving a resolution of up to 100 μm per voxel in an ex-vivo human brain ^8^. However, MRI of human organs at this resolution requires long scan times (100 hrs) and does not achieve the resolution to capture cellular detail. In contrast, multi-beam electron microscopy can provide images of human tissue from cellular to sub-cellular scales ^1^, but cannot capture the large tissue volumes required for whole human organs.

Synchrotron X-ray tomography (sCT) is a promising approach to image whole human organs at cellular detail ^9,10^. X-rays are intrinsically suited to imaging different length scales due to their high penetration and short wavelength. Traditionally, X-ray tomography has provided multiscale 3D imaging for samples above 200 mm in diameter down to a voxel size of 10 μm. To image individual cells requires 1 μm voxels, achievable in large objects only through local tomography, which previously has only successfully been applied to objects with high attenuation differences, such as those found in bones or fossilised remains ^11–13^. There are no established sCT techniques that achieve cellular resolution in large soft tissues such as human organs. Phase-contrast based sCT ^12,14,15^, specifically propagation-based imaging ^16^, can theoretically be applied to image large, soft tissues. However, the X-ray beam coherence and X-ray source brilliance necessary to resolve cellular detail whilst penetrating into samples larger than a few millimetres, was not available at any synchrotron facility worldwide.

Recently, the first high-energy, 4^th^ generation, synchrotron source at the European Synchrotron Radiation Facility (ESRF), named the Extremely Brilliant Source (EBS), has provided the beam coherence required to resolve faint density contrasts at high-resolution whilst achieving one-hundred-fold increase in brilliance compared to its predecessors. We have leveraged the latest iteration of the ESRF-EBS, using the test beamline BM05, to develop a technique termed hierarchical phase-contrast tomography (HiP-CT). We demonstrate the utility of HiP-CT to image intact human tissues across length-scales, spanning whole organs down to cellular resolution in volumes of interest (VOI). We apply HiP-CT to image a series of intact human organs and provide quantitative and morphometric insights into healthy human kidney and lungs. We additionally apply HiP-CT to a contemporary biomedical problem by characterising changes to the architecture of lungs from a patient with fatal coronavirus disease-2019 (COVID-19).

## RESULTS

### A HiP-CT pipeline providing multiscale 3D imaging of large intact soft tissue samples from whole organ to cellular resolution

The overall procedure for the preparation, scanning and reconstruction of intact human organs is shown in **Figure 1A**(see **Online Methods** for detailed steps). Briefly, organs are fixed in formalin solution, partial dehydration in an ethanol series and then physically stabilised for scanning in polyethylene terephthalate jars (**Figure 1B**) using preparations of agar-agar gel. In this configuration, the organs can be mounted or removed between HiP-CT scans at selected resolutions, beginning at 25 μm per voxel for whole organ imaging followed by capturing VOIs at 6 μm per voxel and 1.3-2.5 μm per voxel to achieve cellular discrimination (see **Supplementary Information** for VOI selection template). This configuration is largely automated and can be adjusted to increase scanning speed for smaller organs whilst accommodating different fields of view for high resolution imaging. Beam configuration schematics are provided in **Supplementary Information** and scanning parameters for each sample imaged are detailed in **Supplementary Data 1**. Our experimental procedure allows users to select high-resolution VOIs with spatial context provided by the lower resolution scans and enables systematic or feature-of-interest-based sampling. The registration of images at each resolution is performed via a manually applied rigid transformation (**Supplementary Video 1**).

**Figure 1.**
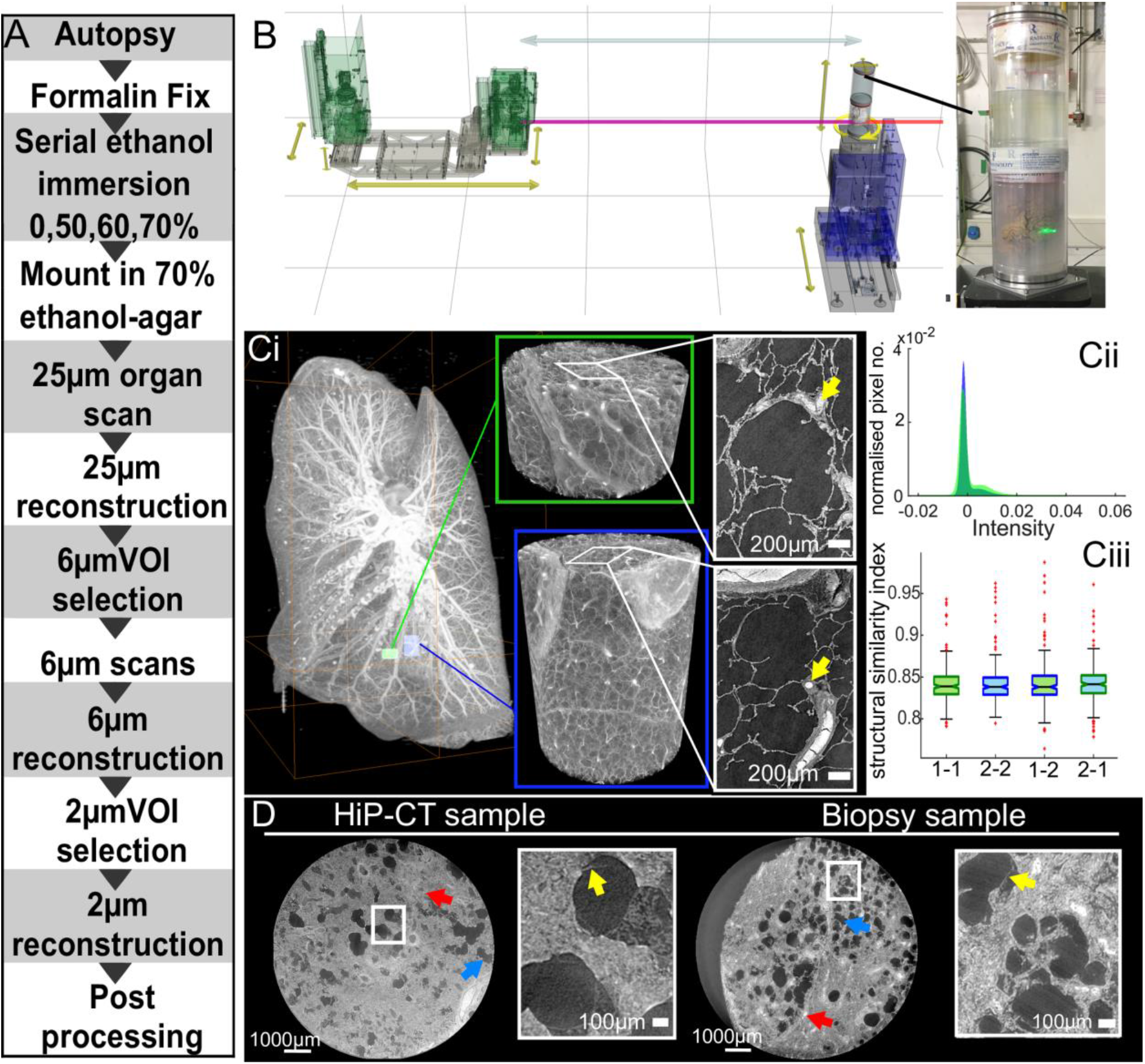
A HiP-CT pipeline for multiscale 3D imaging from whole organ to cellular resolution within large intact soft tissue samples. **A**) Flow chart of HiP-CT human organ sample preparation and imaging. **B**) Experimental configuration, yellow arrows denote possible stage movements, and grey arrow shows propagation distance. The red line indicates the beam. There are two different optics (the two green structures). These provide the range of voxel sizes: 6.5-25 μm is provided by the left hand optic – dzoom optic, and 6-1.3 μm per voxel by the right hand optic-the zoom optic. The sample stage with sample jar can be seen on the right of the schematic and a photograph of the intact human brain mounted in the PET jar with the ethanol agar stabilization is shown in the inset. **Ci**) Maximum intensity projection of whole human lung with two randomly selected VOIs imaged at 2.45 μm voxel resolution shown in green (VOI1) and blue (VOI2). 3D reconstructions of the two high-resolution VOIs are shown, with 2D slices in the insets. In the 3D high resolution VOIs the fine mesh of pulmonary blood vessels, complex network of pulmonary alveoli and their septations can be seen. Yellow arrows denote occluded capillaries in 2D slices **Cii**) Image stack histograms for the green (VOI1) and blue (VOI2) high-resolution VOIs respectively (fixed bin width = 0.0001). Intensity distributions are comparable with positive skew (1.82 and 2.68) and kurtosis (6.44 and 11.88) for VOI1 and VOI2 respectively, histogram intersection is 71% ±3% for fixed bin widths range 1×10^−2^-3×10^−4^. **Ciii**) Box whisker plot showing the structural similarity index between 200 pairs of 2D slices taken randomly either from within the same VOI (1-1 and 2-2) or from different VOIs (1-2 and 2-1; one-way ANOVA; *p* = 0.3217). **D**) Single representative slices of high-resolution scans from a HiP-CT image of an intact whole human lung lobe and a biopsy taken from the same patient’s contralateral lung. Both VOIs are captured from the upper peripheral region of each upper lung lobe. In HiP-CT images, fine structure of the tissue including the blood capillaries (red arrows) and alveoli (blue arrows) as well as thin alveoli septi (yellow arrows in insets) are depicted.

The HiP-CT scanning procedure adapts two protocols originally developed to image large fossils. The first, the attenuation protocol, normalizes the absorption in the field of view ^17,18^ whereas the second, the accumulation protocol, provides extended dynamic range ^19^. After pre-processing of radiographs to generate high-quality local tomography (see **Online Methods** for further details), image reconstruction was performed using a filtered back-projection algorithm, coupled with a single-distance-phase-retrieval ^20^, as implemented in the software PyHST2 ^21^. Subsequently, ring artefacts were corrected on reconstructed slices ^22^.

The entire pipeline, from obtaining fixed organs to 3D reconstruction of intact human organs scans at multiple resolutions is 2-4 days depending on the size of the organ with scanning time for HiP-CT is significantly faster than other similar techniques: ~16hrs for a whole brain at 25 μm per voxel and ~3.5hrs for a whole kidney at 25 μm per voxel (see Online Methods and Table 1. for further details).

**Table 1.**
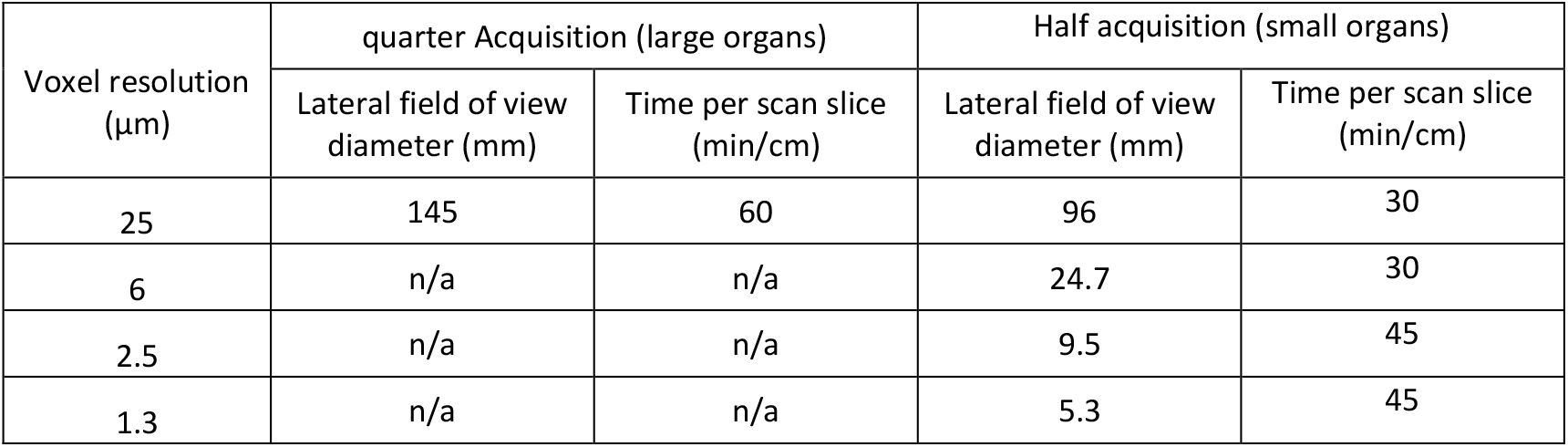
Minimum scan times for quarter and half acquisition modes at resolutions used in this work.

| Voxel resolution ($\mu\text{m}$) | quarter Acquisition (large organs) | | Half acquisition (small organs) | |
| --- | --- | --- | --- | --- |
|  | Lateral field of view diameter (mm) | Time per scan slice (min/cm) | Lateral field of view diameter (mm) | Time per scan slice (min/cm) |
| 25 | 145 | 60 | 96 | 30 |
| 6 | n/a | n/a | 24.7 | 30 |
| 2.5 | n/a | n/a | 9.5 | 45 |
| 1.3 | n/a | n/a | 5.3 | 45 |

To validate the utility of HiP-CT for imaging human organs, we scanned the intact human lung of a 94-year-old female (Donor 1 – clinical details in **Online Methods**) beginning with the whole organ at 25 μm voxel resolution, before randomly selecting VOIs for high-resolution imaging at a 2.5 μm voxel size (**Figure 1Ci**). Demographic and clinical information of all donors is detailed in the **Online Methods**. At low resolution, the 3D architecture of the entire bronchial tree is visible across two lobes of the lung. Within VOIs captured at high resolution, the septations between different alveoli, the functional units of gas exchange in the mammalian lung, and their surrounding blood capillaries can be visualised in two-dimensional (2D) slices and 3D reconstructions (see **Supplementary Video 2**). To assess the consistency in quality of high-resolution HiP-CT imaging at different tissue depths and distances from rotational centre of the intact organ, we analysed intensity distributions across randomly selected 2.5 μm voxel VOIs. We found the image intensity histograms had an intersection of 71%± 3%, both distributions showed positive skew, kurtosis and there was minimal difference in mean intensities between different the different VOIs (**Figure 1Cii**). To further quantify and compare image quality, we used an established comparative measure of image quality - the structural similarity index ^23^ to compare 200 randomly selected image pairs between or within high-resolution VOIs (**Figure 1Ciii**). No significant difference in structural similarity index (median values; 0.839-0.841, *p* = 0.88) was observed across compared images, indicating that HiP-CT can achieve high resolution scanning in any region of the intact human lung with consistent quality. The large size of adult human organs is a fundamental imaging challenge: the penetration depth of the imaging photons (X-ray for HiP-CT) and diffusion of reagents (ethanol for HiP-CT) may be limiting factors, thus image quality tends to decrease from small to large specimens. We therefore qualitatively compared high resolution HiP-CT scans of intact human lung and a physically subsampled biopsy (cylindrical biopsy Ø8.1 mm height 14 mm). The image quality of the two samples was found to be indistinguishable (**Figure 1D**)-the fine structure of the lung tissue including the capillaries (red arrows) and alveoli (blue arrows) are depicted in a histopathologically correct and non-distorted manner with very thin membranes within the alveolae (yellow arrows) being some of the smallest structures (~5 μm thickness) evident in both samples.

Thus HiP-CT is versatile, providing high quality imaging of human lungs across multiple length scales, independent of the region sampled or the size of the tissue imaged.

### HiP-CT enables imaging of organotypic functional units at the scale of whole human organs

Next, we sought to apply HiP-CT to image multiple length scales in a variety of human organs. Intact brain (Donor 2), lung, heart, kidney and spleen (Donor 1) were acquired from two donors (demographic and clinical details in the **Online Methods**), processed and imaged using the HiP-CT pipeline. Images were sequentially acquired at low resolution (25 μm per voxel) to capture the entirety of each organ before high-resolution (6 μm and 1.3 - 2.5 μm per voxel) imaging of selected VOIs spanning each organ’s breadth, (**Supplementary Data 2, Supplementary Videos 1-3**). At 25 μm voxel resolution, macroscopic features of each organ are visible including sulci and gyri of the cerebral cortex (**Figure 2Ai and Aii**), individual lobules of the lung (**Figure 2Bi and Bii**), the four chambers of the heart and associated coronary arteries (**Figure 2Ci and Cii**), the pelvis and calyces of the kidney (**Figure 2Di and Dii**) and the pulpa of the spleen (**Figure 2Ei and Eii**).

**Figure 2.**
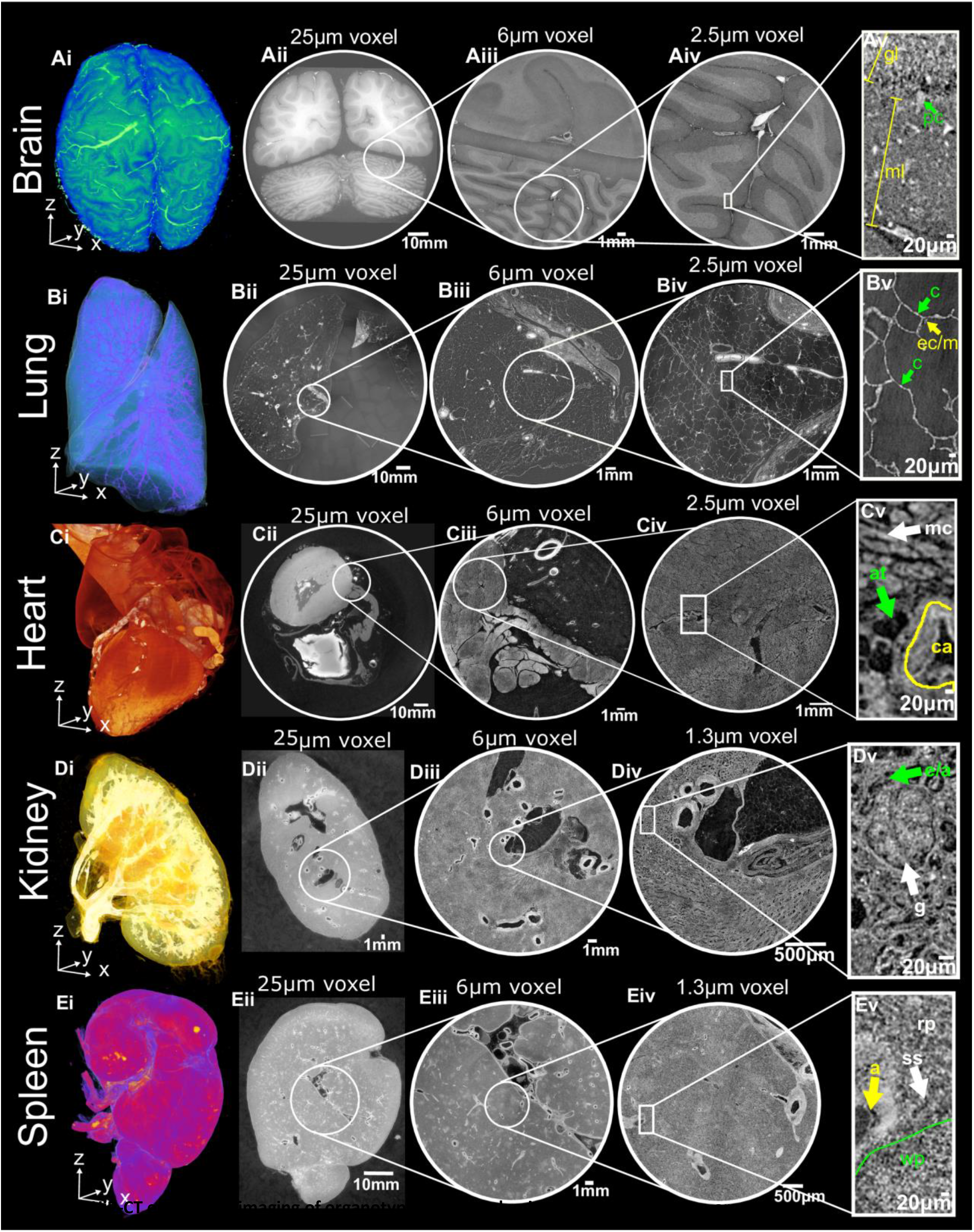
HiP-CT enables 3D imaging of organotypic functional units across intact human organs. HiP-CT of brain (**A**), lung (**B**), heart (**C**) kidney (**D**) spleen (**E**), where for each organ (**i**) shows 3D rendering of the whole organ using the 25 μm per voxel scans. Subsequent 2D slices (**ii-iv**) show the posisitons of the higher-resolution VOIs relative to the previous scan. (**v**) Shows a digital zoom of the highest resolution image with annotations depicting characteristic structural features: in the brain (molecular layer, ml; granular layer, gl; Purkinje cell, pc), in the lung (blood capillary, c; epithelial cell/macrophage, ec/m), in the heart (myocardium, mc; coronary artery, ca; adipose tissue, at), in the kidney (efferent/afferent arteriole, e/a; glomerulus, g) and in the spleen (red pulp, rp; white pulp, wp; arteriole, a; splenic sinus, ss).

Human organ function is driven by the collective activity of organotypic functional units. Each functional unit comprises multiple spatially arranged cell-types that facilitate specialised functions. We used high-resolution HiP-CT to image functional units across the same five human organs (**Figure 2**). In the brain, the architectural layering of the cerebellum and cerebral cortices are visible (**Figure 2Aiii**), and individual Purkinje cells are evident at the base of the cerebellar cortex (**Figure 2Aiv and Av**) which act to coordination cognitive functions ^24^. The termination of the airway into acini is clearly detectable in the lung (**Figure 2Biii**), each containing a cup-shaped alveolus (**Figure 2Biv and Bv**) across which gas exchange occurs. Bundles of cardiac muscle fibres are visible in the heart (**Figure 2Ciii**), consisting of individual cardiomyocytes (**Figure 2Civ and Cv**), which are electrically coupled in a functional syncytium to drive cardiac contraction ^25^. In the kidney, epithelial tubules comprising the nephron are evident (**Figure 2Diii**) which, at their apex, harbour the intricate capillary network of the glomerulus (**Figure 2Div and Dv**), specialised for the filtration of blood. Finally, the organization of red and white pulp in the spleen (**Figure 2Eiii**) are visible, the former of which contains splenic sinuses and the latter contains periarterial lymphoid sheaths (PALS) and lymphoid-rich follicles (**Figure 2Eiv and Ev**). Collectively, these images show that HiP-CT is capable of imaging intact human organs down to the resolution of organotypic functional units and their constituent cells.

### HiP-CT as a tool for mapping functional units (glomeruli) morphology and analysing nephron number

Next, we sought to assess the utility of HiP-CT to obtain physiologically relevant structural information using the human kidney as an example. The functional capacity of the human kidney arises from the collective activity of individual units called nephrons. Each nephron has a glomerulus: a network of blood capillaries which is the site of blood filtration. Therefore, the total number of glomeruli (Nglom) is indicative of nephron endowment within a kidney, and thus its capacity for filtration. No new nephrons are generated through the life course of an individual ^26,27^ and a reduction in N_glom_ is a feature of ageing and chronic kidney disease ^27–33^.

We assessed N_glom_ in a 94-year-old female (clinical information in **Online Methods- Donor1**) by analysing HiP-CT images obtained from their intact autopsied human kidney (**Supplementary Video 3** and **Figure 3Ai**). At 25 μm per voxel, the parenchymal volume was segmented (**Figure 3Aii**) and at 6 μm per voxel, the number of glomeruli was semi-automatically quantified within a representative volume for estimation of N_glom_ (**Figure 3Aiii**). This resulted in a N_glom_ of 310,000 for this individual, which is within the range of N_glom_ in human kidneys found in previous studies using either stereological analysis^32,34,35^, contrast-enhanced MRI^33^ or CT ^32^ and specifically accords well with measures of N_glom_ in older individuals ^35–38^.

**Figure 3.**
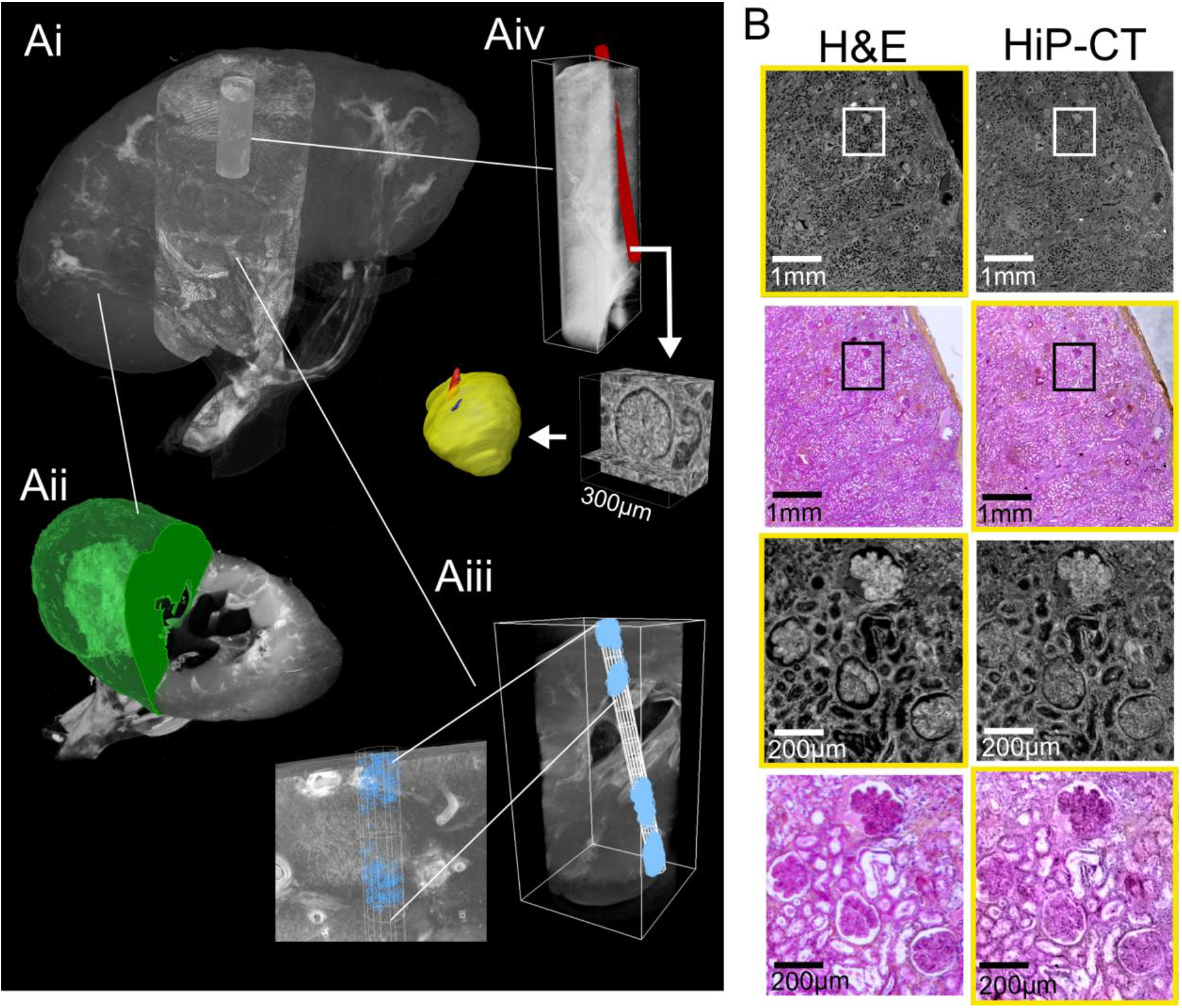
HiP-CT analysis of kidney to measure glomeruli morphology and nephron number. Ai) The three resolution datasets of HiP-CT taken of a human kidney aligned and overlaid. Aii) shows the measurement of the parenchymal volume semi-automatically segmented (green). Aiii) shows the 6μm dataset with the virtual biopsy cylinder in white. 853 glomeruli that were within this cylinder were counted (blue dots) and the parenchymal volume within the cylinder measured. Aiv) the 1.3μm dataset with virtual biopsy (red cylinder) shown. The 13 glomeruli that abutted this cylinder were cropped from the data and segmented (yellow). B) Comparison of HiP-CT with aligned histopathological sections (Haematoxylin and Eosin (H&E) stained) taken after all HiP-CT scanning was finished. The left-hand column shows light micrographs of H&E stained histopathological sections and the right-hand column shows 2D tomograms of HiP-CT, yellow boxes denote images that have been pseudo-coloured.

We also assessed glomerular volume, an important structural determinant of overall kidney health which has been shown to increase in individuals with reduced nephron number, obesity and hypertension ^34,35,39^. To do this, at 1.3 μm per voxel, thirteen glomeruli were manually segmented to calculate V_glom_ (**Figure 3Aiv**) resulting in a value of 5.05 ± 0.09 × 10^−3^ mm, similar to the range of V_glom_ (3-5 × 10^−3^ mm) estimated by stereological and MRI analysis ^33,34,39^. Additionally, the isotropy of HiP-CT enabled the calculation of mean glomerular surface area (1.71 ± 0.4 × 10^5^ μm^2^) and sphericity (0.58 ± 0.09), both of which are unable to be assessed by conventional modalities despite sphericity being a required parameter to approximate V_glom_ from stereology ^30^. Moreover, we matched microscopy of histological sections to the same regions of the kidney imaged with HiP-CT at 1.3 μm per voxel, finding an equivalence of structural detail in the two modalities (**Figure 3B**). Histological results validate the accuracy of glomeruli segmentation and show that sample preparation and high X-ray dose expose during HiP-CT do not cause evident morphological tissue damage (see **Supplementary information** for more details). Therefore, HiP-CT has the potential to quantify functional units and their 3D morphology with histological resolution within intact human kidneys, providing morphometric insights across entire organs.

### HiP-CT demonstrates heterogeneity in parenchymal damage with global changes in alveolar morphology in COVID-19 lungs

We finally sought to apply HiP-CT to a contemporary biomedical problem by examining the structural changes in the lungs of a patient diagnosed with COVID-19, caused by severe acute respiratory syndrome coronavirus 2 (SARS-CoV-2). The major cause of morbidity and mortality in COVID-19 is severe acute respiratory distress syndrome (ARDS), where the efficient transfer of O2 and CO2 between lung alveoli and their surrounding microvasculature is critically compromised ^40^. The clinical course of COVID-19 is well described ^41^ and established histological features of SARS-CoV-2-infected lungs include alveolar inflammation, fibrosis and necrosis ^42^. Recently, these findings were built upon by sCT and 3D reconstruction of millimetre-thick samples from COVID-19 lungs, demonstrating hyaline fibrotic deposits, lymphocytic infiltrates and vasculature occluded by thrombi ^43^.

To assess the utility of HiP-CT to detect lung changes in COVID-19, we imaged an intact upper right lung lobe acquired from the autopsy of a 54-year-old male patient who died from COVID-19-related ARDS (**Figure 4A**). The clinical features of this patient are described in the **Online Methods.** The entire lobe was scanned at a resolution of 25 μm per voxel and reconstructed in 3D (**Figure 4Ai**). Orthogonal 2D slices through the volume of the lung demonstrated high intensity regions in the lung periphery, not observed in SARS-CoV-2-uninfected lung (see **Figure 4Aii** and **Supplementary Video 4**) and consistent with patchy lung consolidation described by conventional clinical radiology in COVID-19 ^44^. Upon higher resolution scanning of VOIs at 6 μm per voxel, 2D slices depicted heterogeneity in the loss of normal alveolar architecture, with particular secondary pulmonary lobules, displaying greater parenchymal deterioration than others (**Figure 4Aiii**). Higher magnification of the more affected secondary pulmonary lobule at 2 μm per voxel (**Figure 4Avi**) demonstrated cavitation of lung parenchyma, alveolar obstruction, (likely from thrombi based on their similar contrast to intravascular blood), thickening of septi between adjacent alveoli and blood capillary occlusion with adjacent cellular infiltrates (likely aggregations of lymphocytes ^42^) (**Supplementary Video 4**). These results indicate that HiP-CT is capable of reproducing the microstructural findings observed in COVID-19 lung biopsies ^42,43^ across large tissue volumes, and in providing access to structures at novel length scales e.g. the secondary pulmonary lobule (**Figure 4B**).

**Figure 4:**
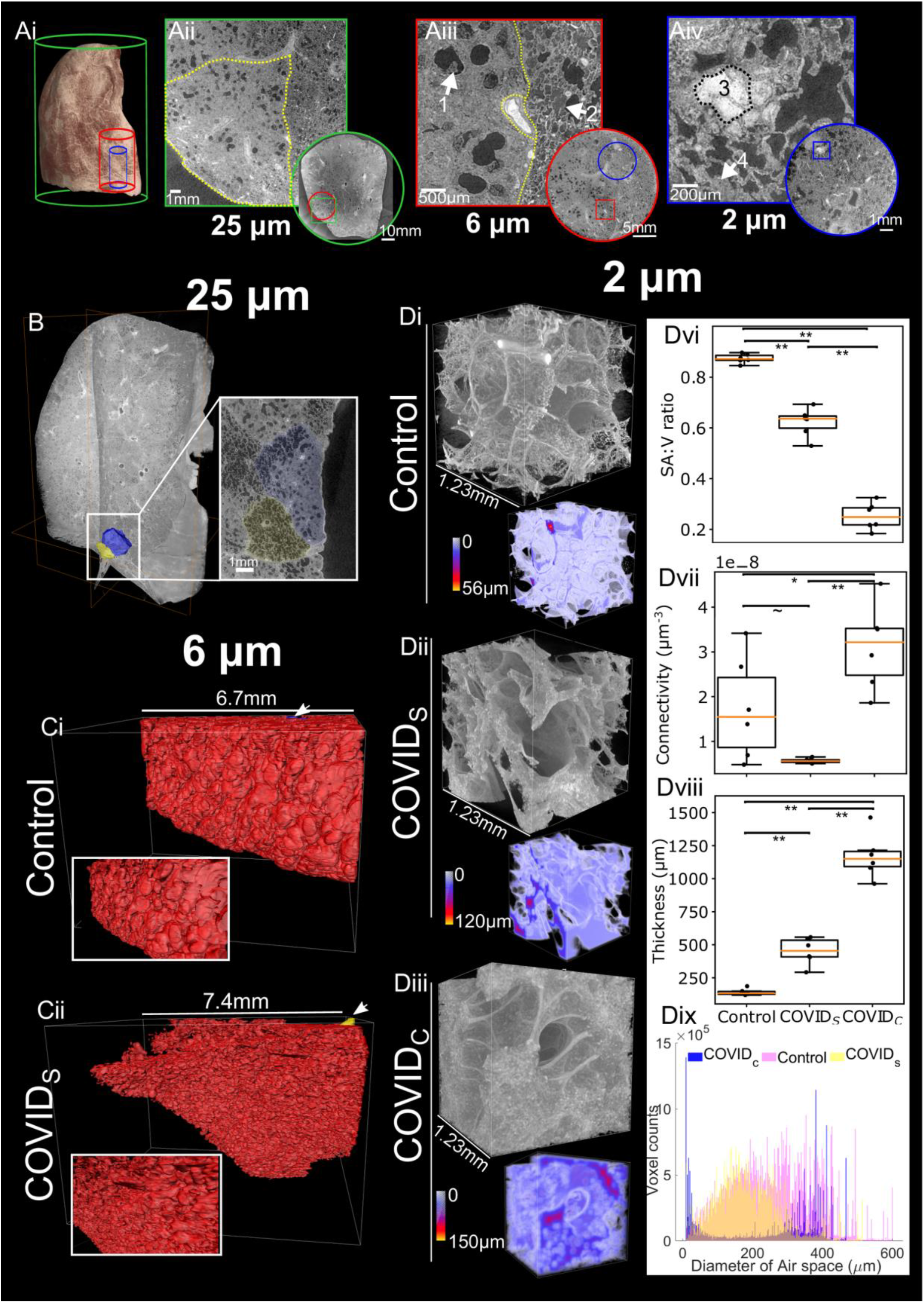
HiP-CT with 3D image analysis and morphometry in the lung of a patient with COVID-19. **Ai**) 3D reconstruction from 25 μm per voxel HiP-CT scanning of the intact upper left lung lobe from the autopsy of a patient decreased from COVID-19-related ARDS. The high-resolution VOIs are shown in red (6 μm per voxel) and blue (2 μm per voxel). **Aii**) At 25 μm per voxel, high intensity regions are observed in the lung periphery. The yellow dashed line delineates a secondary pulmonary lobule. **Aiii**) At 6 μm per voxel, heterogeneity in the lung parenchyma including: (1) dilated alveolar ducts and diffuse loss of alveolar structural organisation; and (2) comparatively well-preserved alveolar structure with some oedematous changes. **Aiv**) At 2 μm per voxel we observed (3) alveolar obstruction, likely representing clotted blood based on its high intensity; and (4) hyaline thickening of alveolar septi. **B**) 3D reconstruction of COVID-19 lung with segmentation of two adjacent secondary pulmonary lobules with differing degrees of parenchymal deterioration. **Ci)** 3D reconstruction of segmented acini structure within the SARS-CoV-2-uninfected (control) lung and **Cii)** the SARS-CoV-2 infected lung. **Di**)-**Diii).** 3D reconstruction of representative VOI at the high resolution (2 μm voxels) for the control, COVID_S_ and COVID_C_ groups respectively. Duplicated volumes show visual representations of the air-tissue interface in COVID-19 lung, where a smaller distance between a voxel of air and a voxel of tissue is coloured blue whereas larger distances are coded yellow. **Div-Dviii**) Boxplots showing quantitative comparisons between COVID_S_, COVID_C_ and control, VOIs. Quantification of mean surface area to volume ratio, airspace connectivity, and mean septal thickness are shown respectively (** indicates p<0.001, * indicates p<0.05 and ~ indicates p=0.08) (calculated by anova with Tukey comparison). **Dix)** Shows the distribution of airspace diameters for all six VOIs in each group modal values for COVID_C_, COVID_S_ and Control = 8.88, 152 and 351 μm respectively.

Given the heterogeneity of parenchymal damage in the COVID-19 lung, we aimed to use HiP-CT to characterise changes of lung architecture in differing regions of the lobe with varying levels of parenchymal deterioration. We segmented two adjacent secondary pulmonary lobules from the periphery of the COVID-19 sample (**Figure 4B**) which showed differing parenchymal deterioration. For the less deteriorated of these two lobules (yellow), we used the higher resolution 6 μm images to segment an acinus (**Figure 4Cii**) and compared this to an acinus segmented from the control lung (**Figure 4Ci**). These images are similar to those obtained by μCT and show the loss of overall surface area in the lung and the alveoli appear smaller and less uniformly shaped. To quantify this difference, we segmented and 3D reconstructed VOIs (n=6 per group) of consolidated lung parenchyma (COVID_C_) and structurally less deteriorated parenchyma (COVID_S_) from the COVID-19 lung and compared these to a control SARS-CoV-2-uninfected control lung (**Figure 4Bi**). 3D quantitative analysis was performed to assess microstructural changes, revealing a significant decrease in surface area to volume ratio (**Figure 4Dvii**) and increase in septal thickness (**Figure 4Dviii**) between control vs COVID_S_, COVID_S_ vs COVID_C_ and control vs COVID_C_ (p<0.001). Airspace connectivity is significantly higher for COVID_C_ than control (p=0.03) and significantly higher than COVID_S_ (p<0.001). The reconstructed VOIs (**Figure 4Di-4Diii**) and the distribution of airway diameters (**Figure 4Dix**), collectively quantify the heterogeneity of lung deterioration in SARS-CoV-2 infection. Thickening of alveoli septae leads to decreased airspace connectivity and a decrease in the modal airway diameter. In consolidated areas, infiltration of loose connective tissue and fluid into the alveoli, leaves only tiny unconnected portions of the alveoli and the large well connected alveoli ducts ventilated. This is evident in the bimodal distribution of airway sizes and high airspace connectivity. Thus, HiP-CT detected regional changes in the architecture and morphology of the COVID-19 lung and allowed large-scale quantification, which can inform our understanding of the pathogenesis of COVID-19-related ARDS in SARS-CoV-2 infection.

## DISCUSSION

Here, we developed HiP-CT, a novel phase-contrast based sCT modality utilising the first high-energy 4^th^ generation synchrotron source. The X-ray beam coherence and brilliance achieved by the ESRF-EBS enabled 3D imaging of multiple intact human organs. Independent of organ size and shape, HiP-CT provides consistent imaging quality through soft tissues, bridging length scales from whole human organs down to individual organotypic functional units and specialised cells. At 1.3 μm per voxel, the current maximal resolution of HiP-CT is over 70-fold higher than state-of-the-art *ex vivo* MRI ^8^ and is comparable to conventional light microscopy. Intact human organs can be processed and multiscale imaged by HiP-CT within days, compared to optical clearing methods for human organs that can take up to 3 months ^7^. We leverage these advantages for multiscale and quantitative 3D analyses of human organ structure, including the estimation of total number, volume, surface area and sphericity of glomeruli in human kidney and the identification of regional changes to the architecture of the air-tissue interface and alveolar morphology in the lung of a donor with confirmed pulmonary COVID-19 disease.

The spatial mapping of human organs is a major challenge of biology. International consortia to map the entire human body and its organs include the Human Cell Atlas and the Human BioMolecular Atlas Program ^45,46^. Such consortia have generated invaluable and openly accessible data of transcriptional states and the cellular and molecular composition of human tissues^47^. To integrate HiP-CT into such efforts, a future challenge is the introduction of labelling methods to capture gene or protein expression at the organ-scale, necessitating the development of suitable radio-fluorescent labels^48^ and characterisation of novel methods for efficient penetration ^7,49,50^ of these labels through large tissue volumes. Another challenge is the computational power and storage required to handle the magnitude of data generated by HiP-CT. A single VOI through the depth of the lung captured at 6 μm per voxel amasses ~600GB. Emerging cloud-based frameworks^51,52^ could improve the speed of image processing and analysis and facilitate the open accessibility of HiP-CT, providing a means to complement global efforts to map the human body at cellular resolution.

HiP-CT has considerable translational potential for biomedical applications, which we demonstrated by 3D imaging of intact SARS-CoV-2-infected lung. In addition to reproducing the histopathological hallmarks of COVID-19-related ARDS, HiP-CT revealed an unprecedented regional heterogeneity in parenchymal damage. This conveys the main disadvantage of physical subsampling by biopsy; the mainstay of histological investigation for many human diseases, which is often used to infer organ-scale pathophysiology from a tiny proportion of total volume of the diseased organ. This sampling bias is surmounted by HiP-CT due to its flexibility in the scale of tissue volume imageable. Moreover, quantitative analysis of preserved volumes of lung parenchyma demonstrated changes in the architecture of the tissue-air interface and abnormalities in alveolar morphology. These results align with clinicopathological observations of an increased volume of ventilated air that does not participate in gas exchange in COVID-19-related ARDS ^40,42^, a phenomenon that has predominantly been attributed to thrombus or embolism within the lung vasculature. Further HiP-CT investigation of COVID-19-related ARDS requires refinement of tools for unbiased feature extraction from scans. We have begun to implement such methods, for example, by training and validating a 3D convolutional neural network ^53^ for automated detection of airway, vascular or parenchymal elements in HiP-CT scans and radiomics ^54^ for high-throughput extraction of radiographic features naked to the human eye (**Supplementary Data**). Similar conventional and novel quantitative methods could be integrated into our HiP-CT pipeline, providing novel insights into the secondary consequences of COVID-19 in other organs, or used to investigate the temporal progression of disease or its resolution in organs such as the kidney and brain; both which show evidence of tropism for SARS-CoV-2 ^55,56^.

HiP-CT will evolve alongside advances in synchrotron technology. Here, we present amongst the first results using the BM05 test beamline. Over the coming years, the completion of a new beamline at the ESRF, BM18, is anticipated to capture cellular resolution in volumes several times larger than human organs whilst using lower X-ray doses. Such 4^th^ generation synchrotron sources may herald new possibilities in the life sciences.

## ONLINE METHODS

### Sample details

Control organs were obtained from two bodies donated to the Laboratoire d’Anatomie des Alpes Françaises (LADAF). Dissections were performed respecting current French legislation for body donation. Body donation was based on free consent by the donors antemortem. All dissections respected the memory of the deceased. The post-mortem study was conducted according to QUACS (Quality Appraisal for Cadaveric Studies) scale recommendations ^57^. COVID-19 lung samples were obtained from the Hannover Institute of Pathology at Medizinische Hochschule, Hannover (Ethics vote no. 9022_BO_K_2020).

Donor 1, from which the heart, lung, kidney, and spleen were imaged, was a 94-year-old, 45kg, 140cm female with right sylvian and right cerebellar stroke, cognitive disorders of vascular origin, depressive syndrome, atrial fibrillation and hypertensive heart disease, micro-crystalline arthritis (gout), right lung pneumopathy (3 years before death), cataract of the left eye, squamous cell carcinoma of the skin (left temporal region).

Donor 2, from which the brain was imaged, was a 69-year-old, 40 kg, 145 cm female, with type 2 diabetes, pelvic radiation to treat cancer of the uterus, right colectomy (benign lesion on histopathology), bilateral nephrostomy for acute obstructive renal failure, cystectomy, omentectomy and peritoneal carcinoma with occlusive syndrome.

Donor 3, The entire upper left lobe and a core biopsy from the periphery of the right upper lobe were obtained from a 54-year-old male patient who died from COVID-19, 21 days after hospitalisation. Treatment involved mechanical ventilation. Regarding comorbidities, arterial hypertension and type II diabetes were not diagnosed prior to death.

### Control Organ Autopsy and Organ Dissection (LADAF)

Bodies were embalmed with formalin solution as follows: embalming solutions - 4500 ml of formalin diluted to 1.15% in a solution containing lanolin and 4500 ml of formalin diluted to 1.44% were injected sequentially into the right carotid artery after death. Bodies were then stored at 3.6 °C. All eviscerations were performed in the LADAF, between April and July 2020.

During evisceration, vessels were exposed, and surrounding fat and connective tissue removed. Organs were post-fixed in 4% neutral-buffered formaldehyde at room temperature for ≥72 hours.

### COVID Autopsy and Organ Dissection

All COVID-19 autopsies were carried out according to standard procedures by the Deutsche Gesellschaft für Pathologie and followed regulatory requirements. Autopsies were performed in a room with adequate airflow (>6 air changes per hour of total room volume) using appropriate personal protective equipment (i.e., hazmat suits, boots, goggles, FFP2/3 masks). Organs were eviscerated and immediately fixed in 4% neutral-buffered formaldehyde. Lungs were immediately inflated and fixed by tracheal instillation with 4% neutral-buffered formaldehyde, the trachea was then clamped, and specimens were left for post-fixation in 4% neutral-buffered formaldehyde at room temperature for ≥72 hours before further dissection. Samples were partially dehydrated to 70% ethanol (through four successive concentrations see below) prior to transportation to ESRF.

COVID lung samples were transported to ESRF, Grenoble at ambient temperature in double-sealed containers.

### Organs preservation and degassing

In order to ensure resistance of the samples to the X-ray dose, avoid health risks associated with formalin and to increase the general contrast level; all the organs were equilibrated in a 70% ethanol solution. As the ethanol replaces water in tissues, the density of the water-rich tissues and water-filled cavities decreases, while the density of other tissues remains closer to the original one. This results in improved X-ray contrast, especially visible when using propagation phase contrast imaging.

In order to avoid shrinkage of the organs, the conversion from formalin to 70% ethanol was performed in four successive baths: 50% (volume of solution 4 × volume of organ), 60%, 75% (to reach an equilibrium at 70% taking into account the volumes of the organ and the solution), and finally, 70%. Degassing was performed at each step with the organ immersed and blocked in the respective solution using a diaphragm vacuum pump (Vacuubrand, MV2, 1.9m^3^/h).

Degassing was performed through cycles of vacuum and slow return to atmospheric pressure with increasing duration from 2 minutes to 40 minutes (total time typically 2h). At each cycle, the pumping was stopped when bubbling started to decrease in intensity. In the case of the Donor 1 control lung, after each cycle, the degassed ethanol solution was gently forced into the main conducting air ways with a large syringe in order to keep the morphology of the lung in its original inflated state. This series of pumping cycles aims to remove all free gas in the organ (especially in the cases of lungs), and as much of the dissolved gas in the tissues as possible.

The final degassing step, in the 70% solution was longer (typically 3-4 hours) until no bubbling was observed. Samples were then mounted for the scanning process.

### Organ mounting with Agar-agar ethanol gel preparation

The mounting of the organs for scanning requires delicacy because any remaining bubbles, density inhomogeneities or insufficient sample stabilization would dramatically reduce the quality of the scans. We decided to use 70% ethanol-based gel in order to ensure that no chemical diffusion occurred between the organ and the mounting media. After several tests, the most suitable solution was based on agar-agar gel.

As it is not possible to prepare directly agar-agar with an ethanol solution, the gel was first prepared using demineralized water (20g/L of agar agar). Once gelled, the block of gel was cut into small cubes of 1cm^3^ that were immersed in ethanol at 96%, with a volume ratio ensuring the final concentration at equilibrium of 70% of ethanol (2.96L of ethanol at 96% for 1L of agarose gel). Agarose gel does not shrink when immersed in high concentration ethanol, and it reaches a concentration equilibrium with the solution (upto 24 hours). The equilibrium state can be tested by checking that gel cubes sink (over several dozen seconds) to the bottom of the container after agitation, indicating that their density is very close to the solution. Once equilibrated, the gel cubes still immersed in the solution were degassed using the vacuum pump for approximately one hour, (until no bubbles form).

Approximately a third of the cubes are kept in this degassed equilibrated state (always stored in the solution to avoid drying that would result in higher density and shrinkage). The other two thirds were blended into a homogeneous viscous media (termed the “blended gel”), the consistency of which was adjusted by adding more ethanol at 70% from a prepared solution.

### Organs mounting and degassing

During the exploratory phase of this project, several mounting protocols were tested. The first one (used for the COVID-19 lung-lobe presented in this paper), consisted of using only the agarose gel cubes in 70% ethanol solution to fix the sample in the plastic sealed tube used for the scan. This preparation allowed efficient final degassing but causes unwanted local compression of the soft organs that can affect their structures near the surface (as visible on the **Supplementary Video 4**). In some cases, it also leads to the agar blocks being visible in the reconstructed scans, as the top surface of the agar-agar gel may be of slightly higher density than the bulk, if too much time elapses between the gelation and the cutting (partial dehydration of the surface).

In order to have a more homogeneous mounting media, we developed a three-step protocol: the bottom of the cylindrical container was filled with gel cubes in a few centimetres of blended gel. These cubes ensure that the organs cannot contact the base of the container during mounting and contribute to preventing eventual rotations of the complete mounting media in the container. Then the tube was half-filled with blended gel, and the organ was carefully immersed and set in the desired position (with special care to avoid bubbles). The tube was filled again with blended gel up to a few centimetres above the organ. The whole mounting was then degassed once again, with special attention to ensure that the gel did not inflate too much due to bubbles trapped during the process. Once degassed for 1h, the mounted organ was covered with gel cubes that were pushed in the top layer of the gel to ensure solid enough fixation of the organ. The whole mounting was degassed a last time for few minutes, then the container was sealed for scanning.

### Specific sample holder and mounting

We used specially designed sample holders for the scanning process. They were designed to ensure safety in case of a leak of the ethanol via a double sealing, to be mechanically stable and resistant to high X-ray dose. Two samples can be mounted one over the other for longer scan automation. Alternatively, only one organ can be installed, and the equivalent organ container filled with 70% ethanol solution is installed in the second place for beam reference measurements in case of local tomography scans (**Figure 1B**).

### Multi-resolution scanning protocol

One organ in its sealed container was installed in the bottom holder, and the equivalent container filled with 70% ethanol gel was installed in the upper holder (**Figure 1B**). This second container was used for acquisition of beam references with the exact same geometry and scanning parameters as the organ scan (**Figure 1B**). By using a reference scan to perform the flatfield correction of the radiographs of the organ scan, most of the low frequency effects of local tomography are directly corrected, leading to normalized intensities of the projections even in case of off-center local tomography scans.

This approach also makes it possible to improve the dynamic range of the detector by avoiding direct exposure to the beam. The sample and its mounting media act as filters. This protocol, we term the “attenuation protocol”, is derived from an approach that was originally developed to scan highly absorbing fossils, first in monochromatic beam ^17^, and later in polychromatic beam ^18^. In order to maximize the dynamic level without using a dedicated beam profiler for scanning organs at 25μm, the setup was put slightly off axis to benefit from the natural horizontal profile of the beam. Using this approach, it was possible to reduce the flux on the border of the sample (where the absorption is lower), and to use the maximum flux in the largest dimension of the sample. These are all adaptations from previous attenuation protocols from fossils studies to biological samples but the application to off-axis scans is entirely novel. The addition of the second container and equivalent scan geometry for adaptive references during rotation enables scanning with any position and resolution in our whole organ samples.

In addition to correction of the local tomography effect, our scan protocol allowed us to reach a high dynamic level by avoiding direct exposure of the detector. The saturation level of the detector can then be adapted to the less absorbing part of the sample instead of to the direct beam intensity. As the general absorption contrast is dominated by the cylindrical shape and the organs are close in density to the ethanol solution, this approach also efficiently removes any beam hardening effect, or differential phase contrast effect in case of off-axis scans. Finally, in order to ensure a high dynamic level, the detector is used in accumulation ^19^, with typically 10 and 5 sub-frames for the 25 μm and 6.5μm scans respectively. The 2.5 μm and 1.3 μm scans are done without accumulation in order to limit the X-ray dose.

All the scans were performed on the beamline BM05 of the ESRF using two different optics. The dzoom optic covered pixel sizes from 25 μm to 6.5 μm (**Figure 1B and Supplementary Figure 1A**). The zoom optic covered from 6.2 μm to 1.4 μm (**Figure 1B, Supplementary Figure 1B**). Both optics were mounted with PCO edge 4.2 CLHS cameras.

Two acquisition modes were used depending on the size of the organ samples – half and quarter acquisitions. Most scans were performed in half-acquisition mode with the centre of rotation on the right side of the field of view (moved by typically 900 pixels) to obtain a field of view of 3800 pixels with 6000 projections. For the largest organs, the complete scans at 25μm were performed using a quarter acquisition protocol based on two scans (one half-acquisition + one annular scan) of 9990 projections each. Once concatenated, the reconstructed field of view is 6000 pixels. In order to cover complete organs or to scan large columns in local tomography, scans were performed with automatic series along the z axis. The typical sampling step was 2.2mm vertically for a corresponding beam size of 2.6 mm (overlap of 18%). Nevertheless, some organs scanned at the beginning of the project have been scanned with larger steps (up to 3.6 mm), and larger overlapping (up to 50%).

**Supplementary Figure 2** shows the motor limits and how VOIs were selected in the half and quarter acquisition modes.

All the details of the scanning parameters for each scan of each sample are presented in the **Supplementary data 1**. Details the lateral fields of view and fastest possible scan times currently available for the different resolutions and acquisition modes are provided in **Table 1.** Based on these times the brain supplied by Donor 2 can be scanned at full 25 μm resolution in ~16hrs (16cm); and the smallest diameter organ – the kidney (7cm) in ~3.5hrs.

### Tomographic data reconstruction protocol

After pre-processing, the tomographic reconstruction was performed using the filtered back-projection algorithm, coupled with a single distance phase retrieval ^20^, and with a 2D unsharp mask on the projections, as implemented in the ESRF inhouse software PyHST2 ^21^. All the sub-volumes are converted into 16 bits, and vertically concatenated. The remaining ring artefacts were corrected on the reconstructed slices using an inhouse Matlab system derived from Lyckegaard et. al ^22^. For the most difficult cases, a final correction of the horizontal stripes was performed on the reconstructed volumes after vertical reslicing. Nevertheless, this step was typically not necessary when the stripes had already been corrected on the concatenated radiographs.

All the codes used for this processing are either available in supplementary information or were already available on open-source libraries through previous publications^21,22^. They have been developed specifically for HiP-CT by P. Tafforeau and are provided in their native status.

### Histology (haematoxylin and Eosin)

After HiP-CT scanning was complete the kidney was removed from the mounting jar. The coronal location of the kidney that approximately aligned with HiP-CT high resolution image columns were manually identified. Coronal sections were placed into large cassettes to follow the standard tissue treatment procedure: dehydration through a series of graded ethanol baths, paraffin embedding and microtome sections of paraffin blocks. All slides were routinely stained with haematoxylin and eosin. Slides were visually inspected to find more precisely aligned images to high resolution HiP-CT.

In addition to performing H&E on the HiP-CT imaged kidney in the portions of the sample that received the highest X-ray dose (**Figure 3B** and **Supplementary Figure 3)**, staining was also performed on the COVID lung biopsy sample shown in **Figure 1D** and on healthy control lung biopsies. **Supplementary Figure 4** shows H&E staining as well as anti-CD31, anti-thyroid transcription factor 1 (TTF-1) and anti-fibrin immunohistochemical staining on the lung biopsy samples. As these lung samples were biopsies, they did not receive the high doses the kidney was subjected to (see **Supplementary information** for discussion and estimation of does exposure) however these sample underwent similar preparation and were exposed to the ESRF-EBS on BM05.

### Image Analysis

The final volume renderings are performed using different 3D software packages, notably VGStudioMax 3.2.4 (Volume Graphics, Heidelberg, Germany), Amira v2019.6 and Dragonfly 2020.2.0

### Image quality comparison (Figure 1Cii and Ciii)

Image quality comparison was performed using the structural similarity index ^23^ implemented in Matlab 2020b (ssim function).

Equally sized subvolumes from the two columns were used in their original 32-bit form (999 slices in each case). Histograms of the two subvolumes were calculated with fixed bin width of 0.001. Skew, kurtosis and mean pixel intensity were calculated via Pyradiomics v3.0.^54^.

Structural similarity index was calculated between pairs of xy slices from the two subvolumes. Two images were randomly selected (by slice number); either both images were from the same subvolume (1-1) and (2-2) or the images were from different subvolumes (1-2) and (2-1). 200 pairs of images were analysed for each group. One-way anova of the output was performed in Matlab 2020b using the anova1 function.

### Kidney Glomerulus analysis

Using the 25 μm per voxel scans of the whole kidney, the parenchyma was semi-automatically segmented in Amirav2019.6. After manually aligning all three resolution datasets, two cylinders, one through each of the 6μm and 1.3μm columns, were defined perpendicular to the kidney surface. The cylinders had radii, lengths, and volumes of (1300μm, 40,811μm and 2.17×10^11^μm^3^) and (203μm, 9814μm and 1.3×10^9^μm^3^) for the 6μm and 1.3μm datasets respectively. These cylinders were used as virtual biopsies. In the 6μm case any glomeruli that fell within the cylinder were counted (blue dots in **Figure 3Aiii**). In the 1.3μm dataset any glomeruli that touched the cylinder were manually segmented in Amira v2019.6 in 3D. The volume, surface area and sphericity were calculated using the binary mask from the segmentation in ImageJ using the MorphoLibJ plugin-(analyse regions 3D)^58,59^. For the total number of glomeruli, the volume of parenchyma in the 6μm virtual biopsy was found via semi-manual segmentation in Dragonfly 2020.2.0. The number of glomeruli counted in the 6μm virtual biopsy was divided by the volume of parenchyma in the 6μm virtual biopsy and multiplied by the total volume of the parenchyma. Alignment and pseudo colouring of the HiP-CT and H&E histology sections was performed in VGstudioMax 3.2.4 by altering the transparency and colour map of each image.

### Lung microstructure analysis

Manual segmentation of secondary pulmonary lobules was performed in Amira 2019.6 using the 25 μm COVID-19 upper right lung lobe. The data was binned to 50 μm prior to segmentation to reduce computational load. Acini segmentation was performed using a randomly selected column of 6 μm in the control lung and a region of the COVID-19 lung selected by pulmonary radiologist as relatively spared. Regions of the image that contained a clear terminal bronchiole were cropped and segmentation was performed semi-manually: first tissue air boundaries were enhanced using an unsharp mask and a Canny edge filter in ImageJ ^58^, enhanced images were then transferred to Amria2019.6 and a 3D magic wand tool was used segment the 3D airspace. For microstructural analysis, six 250×250×250pxl subvolumes from the 2.2 μm datasets of the COVID-19 and control lung, were chosen, (randomly in the case of the control lung, and in the COVID-19 case, randomly within larger areas designated by pulmonary histopathologists as structurally preserved or consolidated i.e. from areas where alveoli morphology was identifiable verses areas where there was widespread consolidation of the parenchyma). Binary images separating airspace from alveoli septea and small vessels were produced in ImageJ by thresholding the images used one two methods: ‘triangle’ method for Control and COVID_S_ and ‘Yen’ method for COVID_C_ 58. A 3D Chamfer distance from the MorpholibJ plugin ^59^ was used (using Svensson 3,4,5,7) to create the distance maps (Figure 4Di-iii). The BoneJ plugin (V2) was used to perform morphological analysis including connectivity, surface area to volume ratio, airway diameter and septal thickness^60,61^. One-way anova with Tukey comparison was used for statistical analysis of microstructural measures.

## Acknowledgements

Sam Bayat (ISERM), Philippe Masson (LADAF) for body donors’ dissections, Harald Reichert (ESRF) and R. Tori for general support of the project, and Clemence Muzelle, Roberto Homs, Christophe Jarnias, Filippo Cianciosi, Philippe Vieux, Phil Cook, Luca Capasso and Alessandro Mirone for their help in the setup developments and improvements. The authors also thank Regina Engelhardt, Annette Muller Brechlin, Christina Petzold and Nicole Kroenke

## Funding

This project has been made possible in part by grants number 2020-225394 from the Chan Zuckerberg Initiative DAF, an advised fund of Silicon Valley Community Foundation, The ESRF - funding proposal md1252, the Royal Academy of Engineering (PDL - CiET1819/10) and the MRC (MR/R025673/1). CW is supported by the MRC Skills Development Fellowship (MR/S007687/1). DDJ and DAL are supported by funding from, Kidney Research UK (Paed_RP_10_2018, IN_012_2019), the Rosetrees Trust (PGS19-2/10174, PhD2020\100012), a Wellcome Trust Investigator Award in Science to DAL (220895/Z/20/Z), the UCL MB/PhD programme and a Child Health Research PhD studentship to DJJ and the National Institute for Health Research Great Ormond Street Hospital Biomedical Research Centre. MA acknowledges the European Consolidator Grant, XHale (ref. no.771883) and the National Institutes of Health (HL94567 and HL134229). JJ acknowledges Wellcome Trust Clinical Research Career Development Fellowship 209553/Z/17/Z and the National Institute for Health Research University College London Hospital Biomedical Research Centre. This work was supported by the German Registry of COVID-19 Autopsies (DeRegCOVID, www.DeRegCOVID.ukaachen.de; supported by the Federal Ministry of Health - ZMVI1-2520COR201), and the Federal Ministry of Education and Research as part of the Network of University Medicine (DEFEAT PANDEMIcs, 01KX2021).

## Notes

### Competing Interest Statement

The authors have declared no competing interest.

