## Supplementary Data 2 for "Multiscale three-dimensional imaging of intact human organs down to the cellular scale using hierarchical phase-contrast tomography"

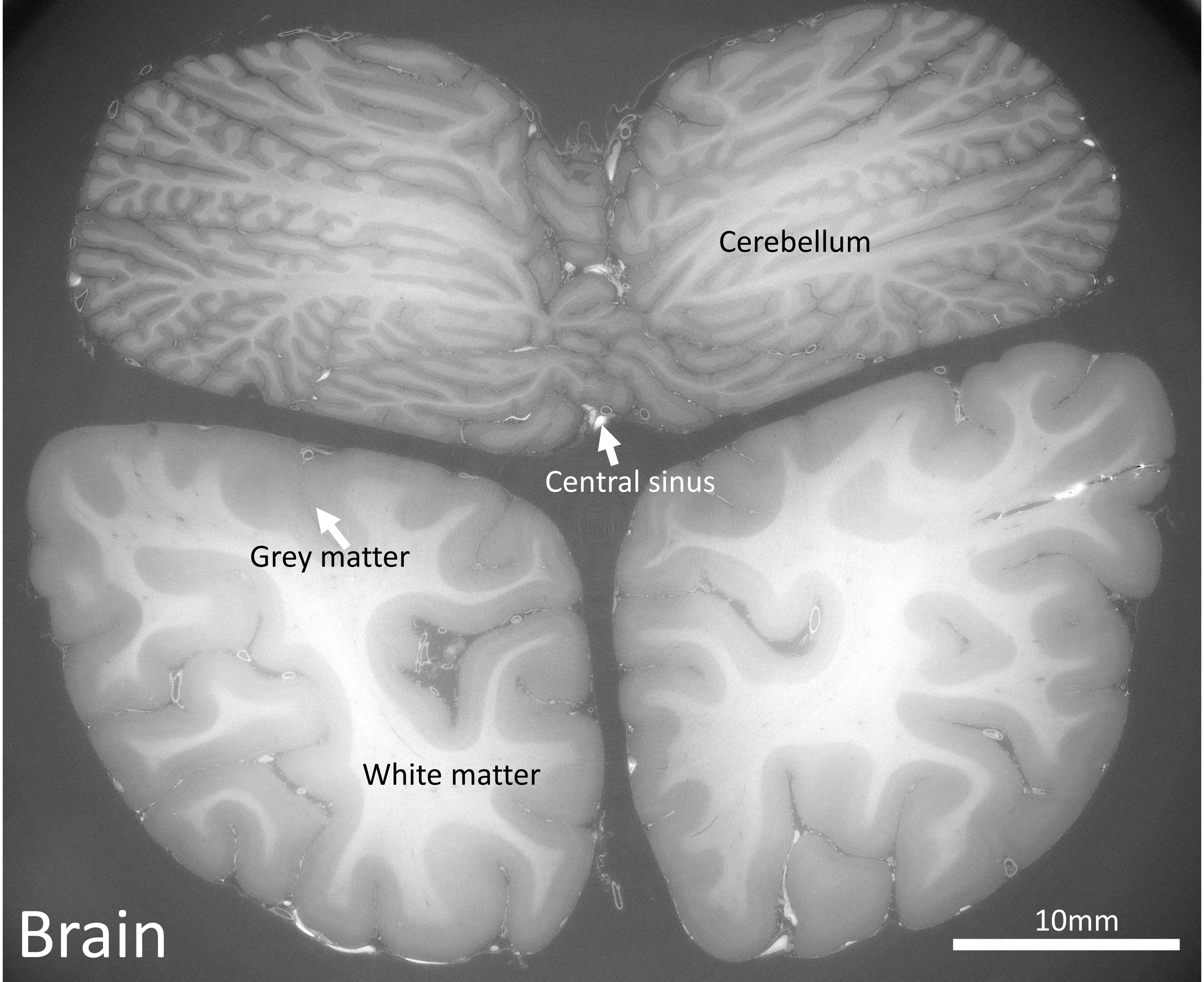
Cerebellum
Central sinus
Grey matter
White matter
Brain
10mm

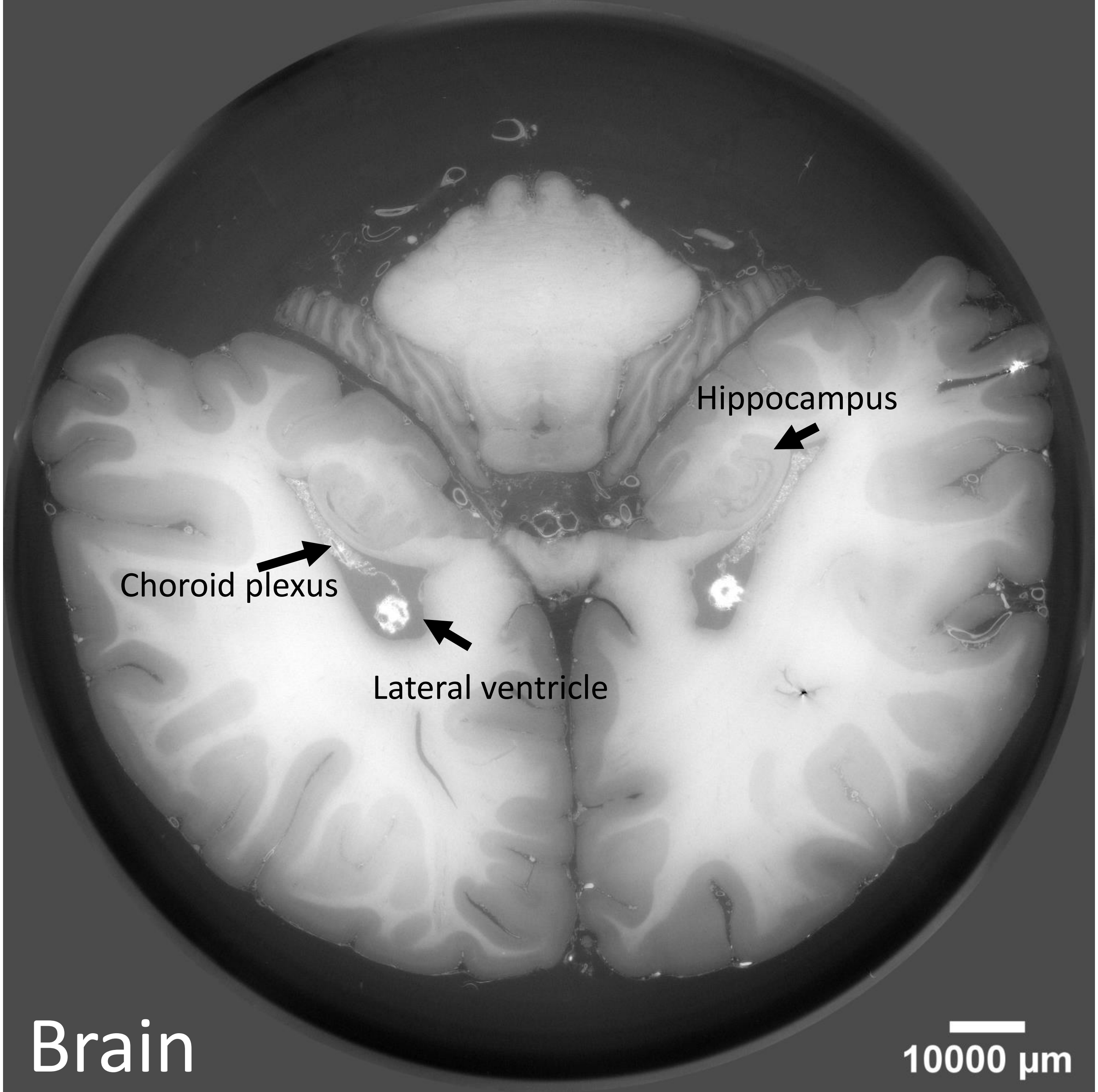
Hippocampus
Choroid plexus
Lateral ventricle
Brain
10000 μm

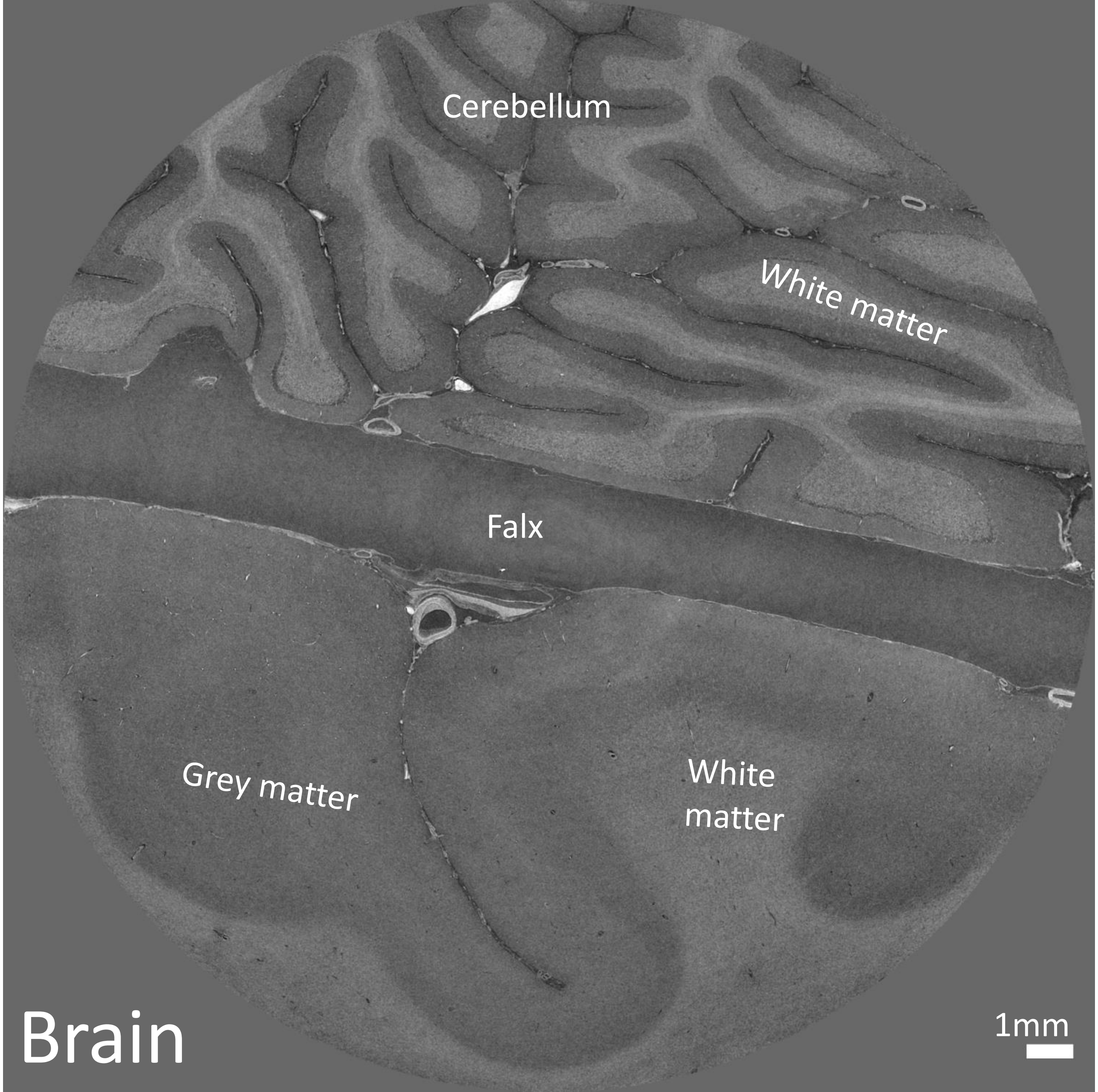
Cerebellum
White matter
Falx
Grey matter
White
matter
Brain
1mm

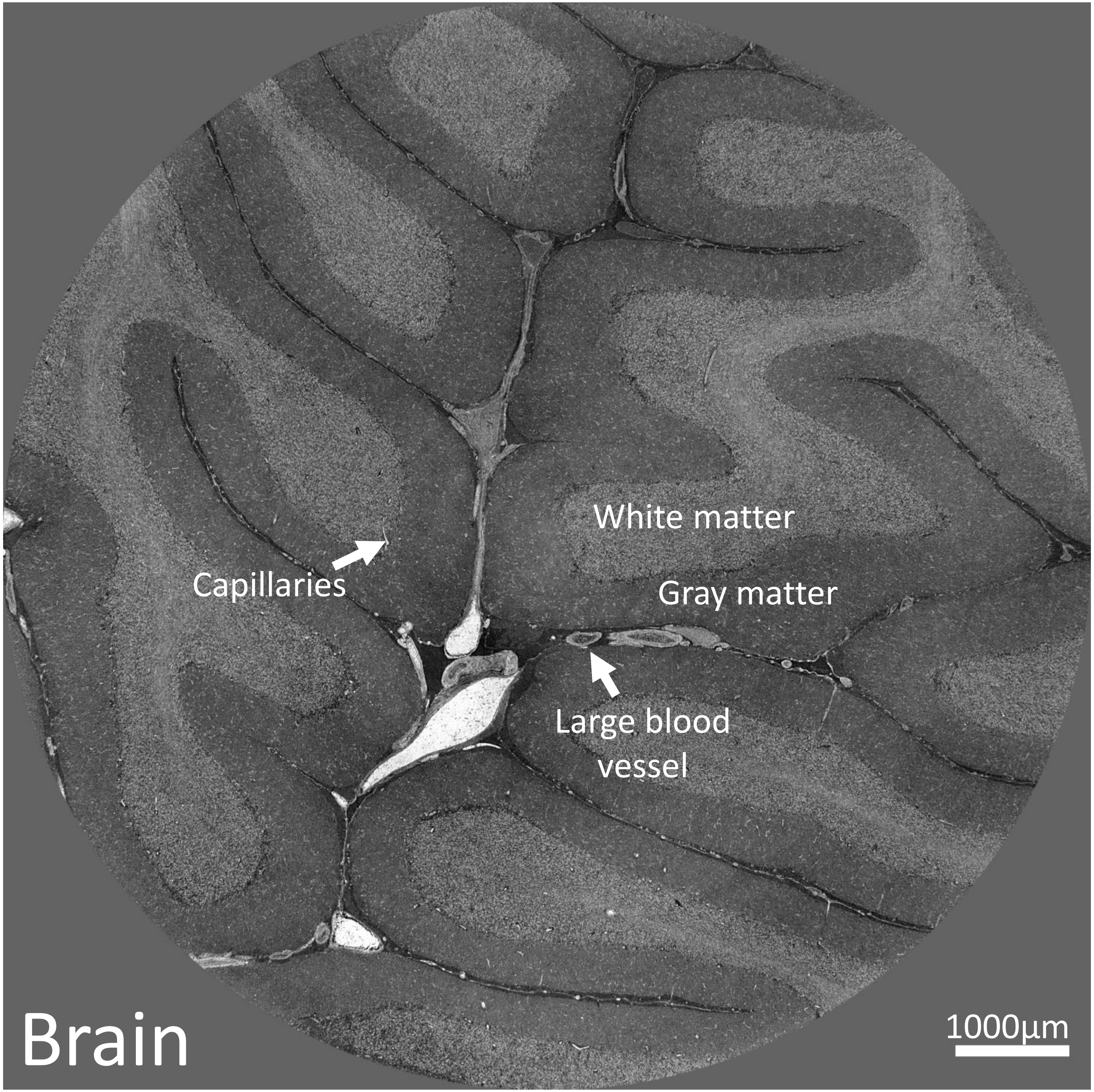
White matter
Capillaries
Gray matter
Large blood vessel
Brain
1000µm

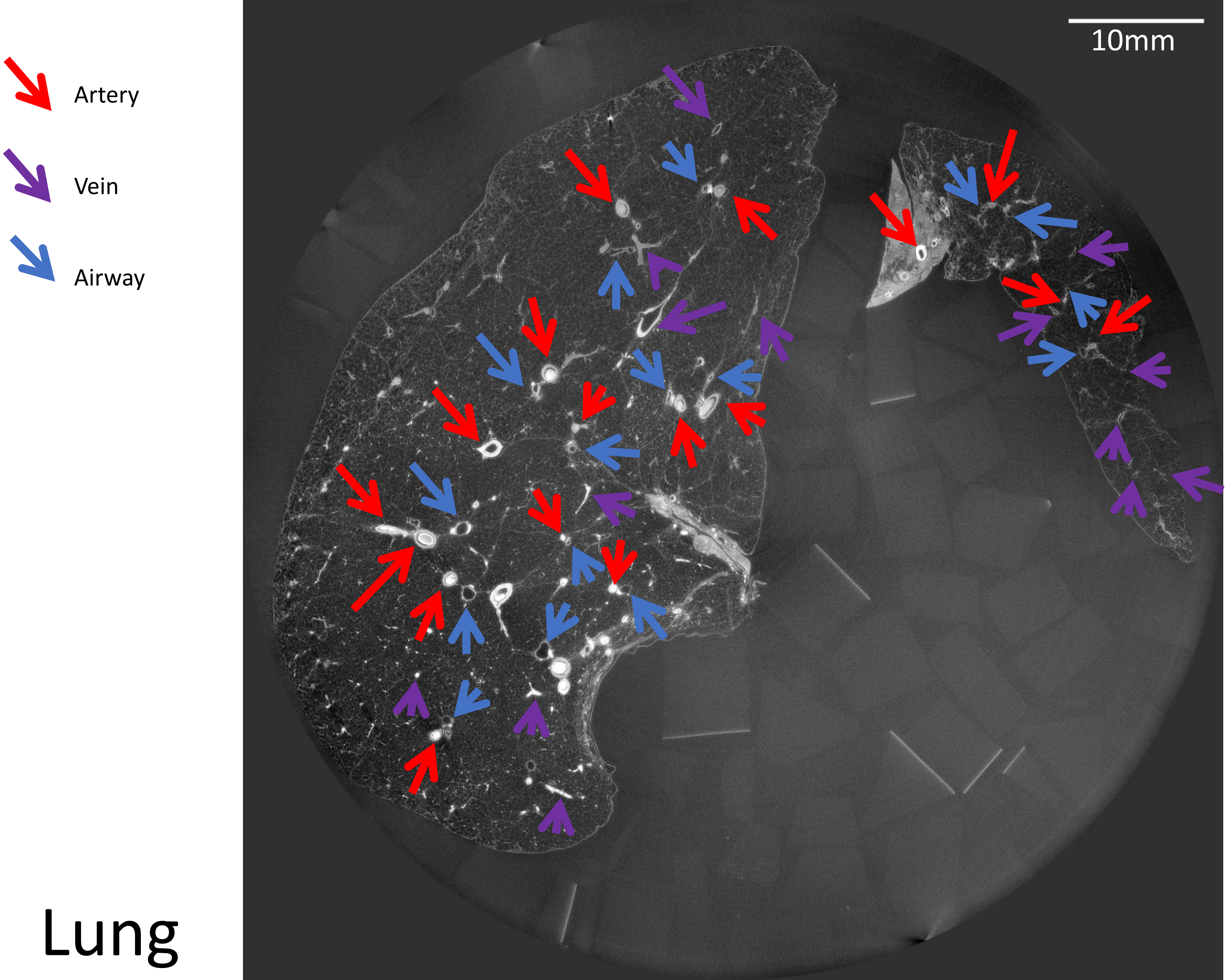


### Lung

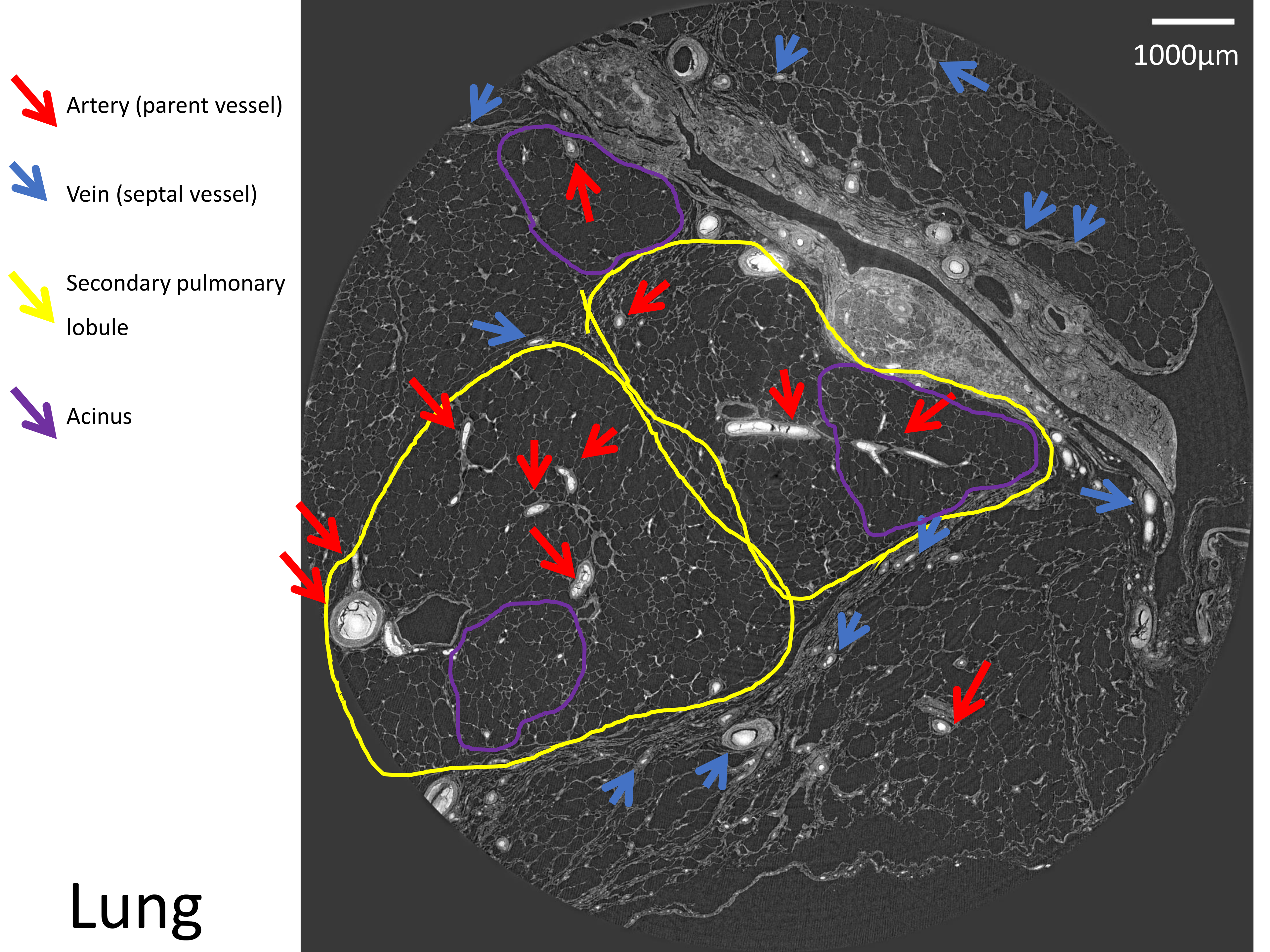
Artery (parent vessel)
Vein (septal vessel)
Secondary pulmonary lobule
Acinus
1000µm
Lung

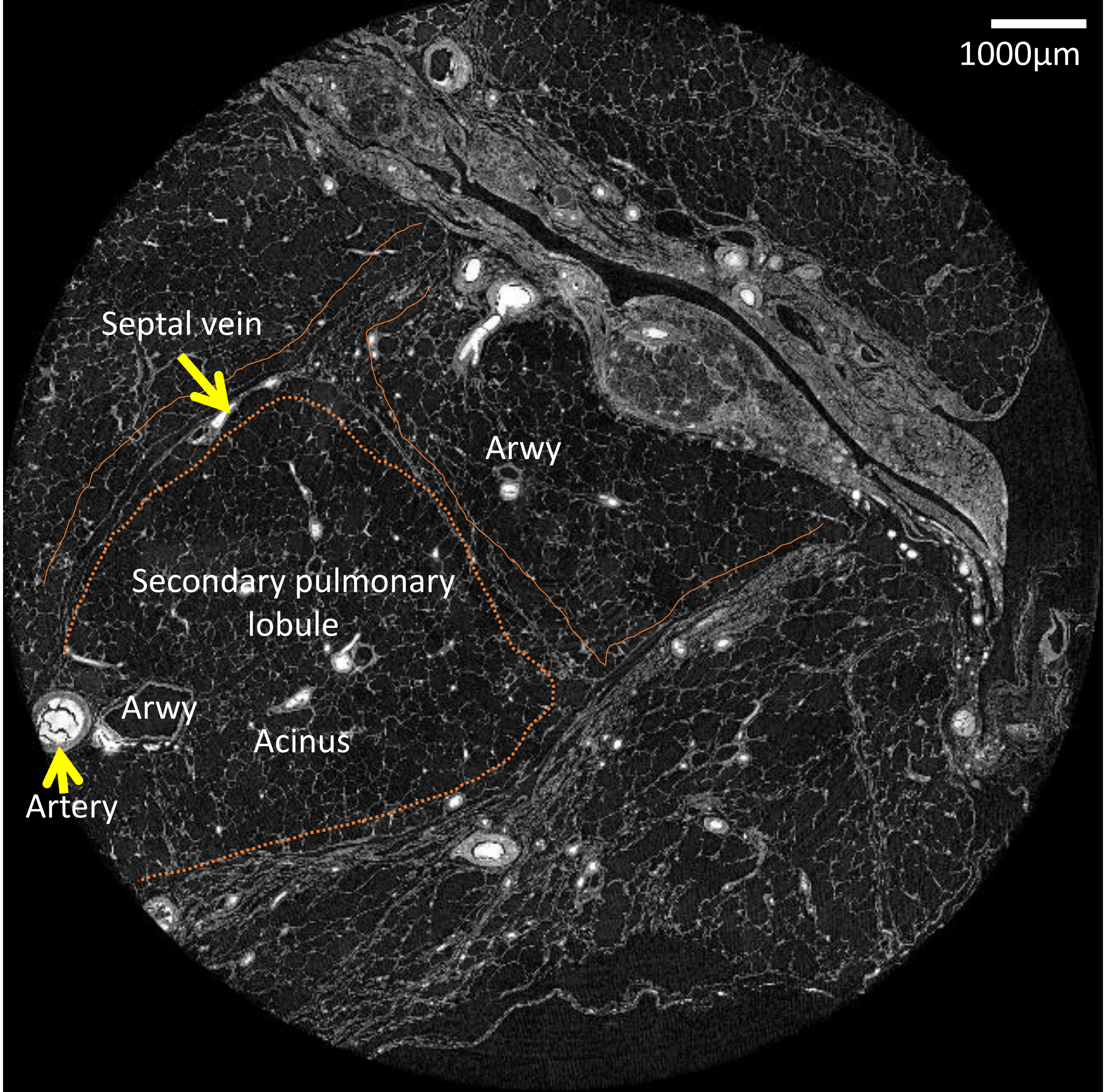
1000µm
Septal vein
Arwy
Secondary pulmonary
lobule
Arwy
Acinus
Artery

### Lung

Capillaries

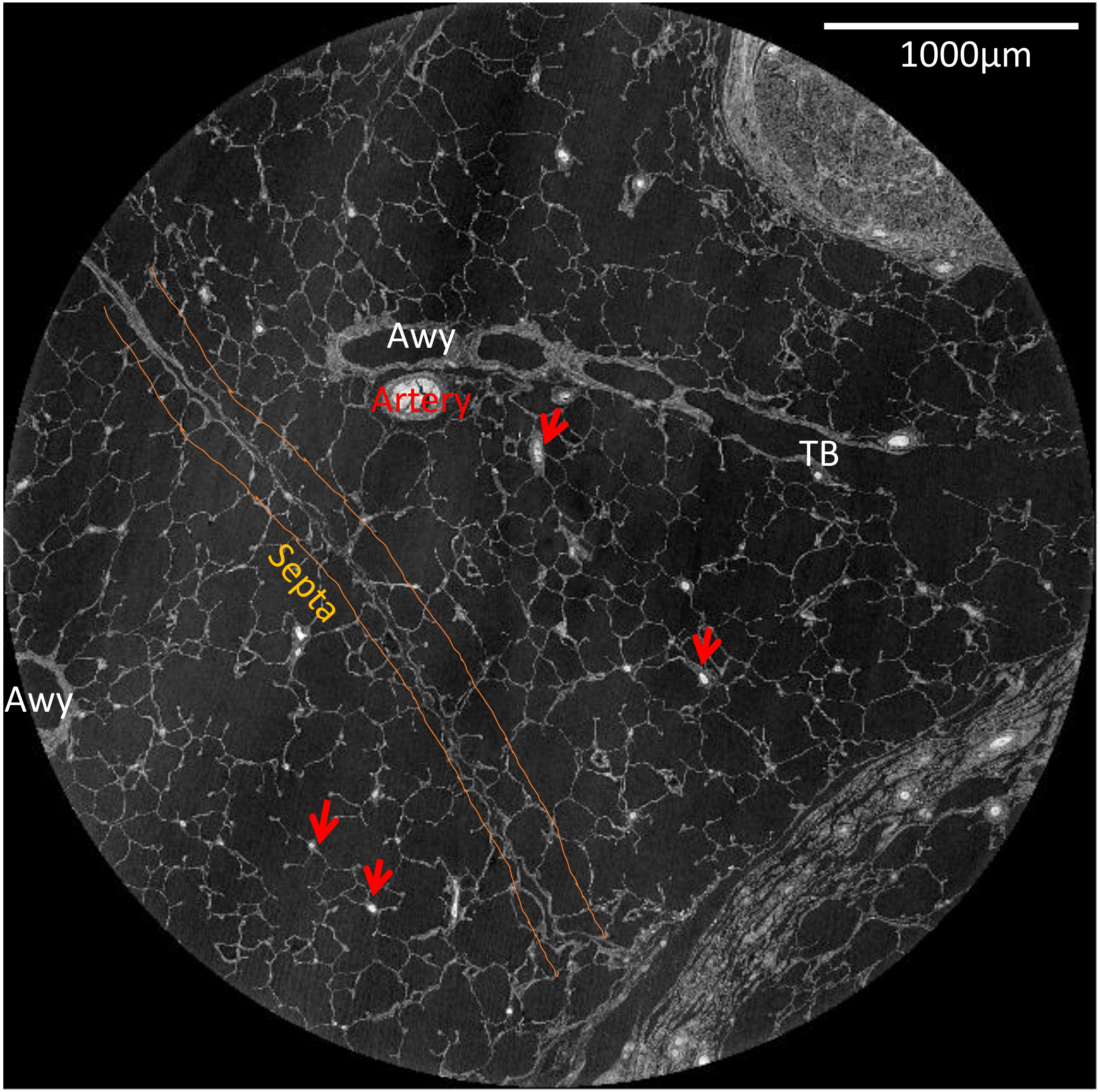


### Lung

Single capillaries/ endothelial cells

Alveolar epithelium cells (type I or II) / macrophages

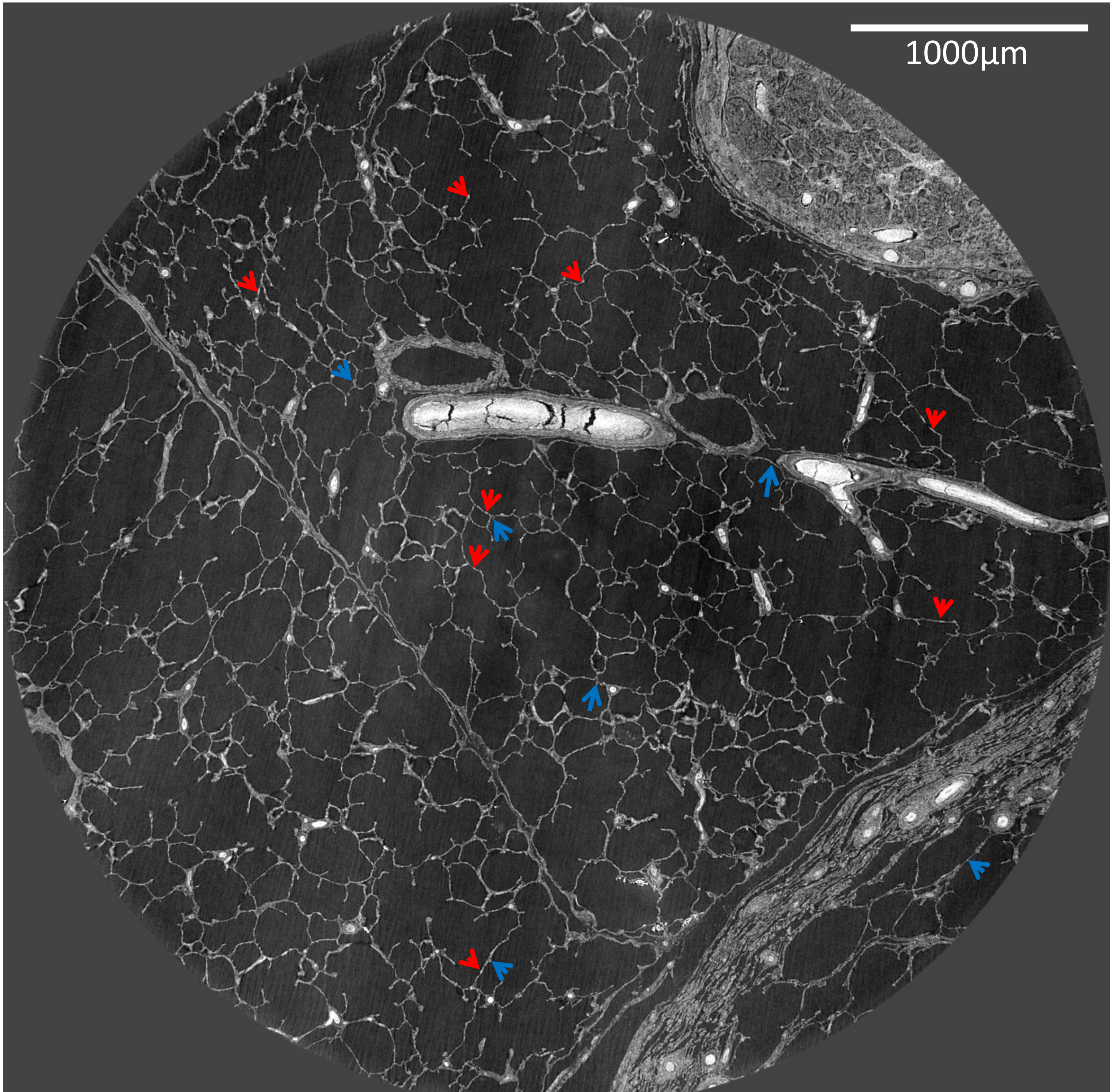

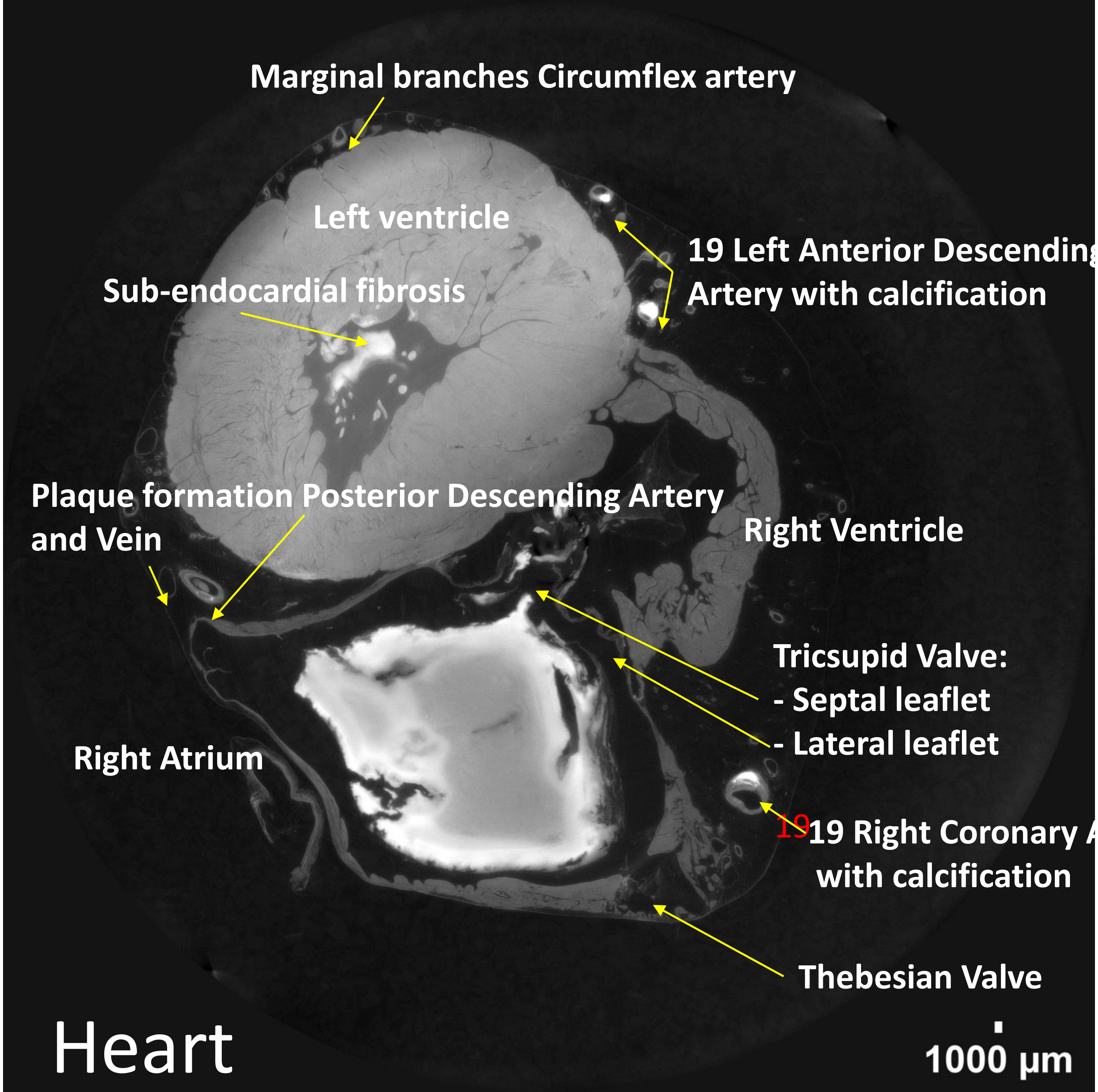
Marginal branches Circumflex artery
Left ventricle
Sub-endocardial fibrosis
19 Left Anterior Descending
Artery with calcification
Plaque formation Posterior Descending Artery
and Vein
Right Ventricle
Tricsupid Valve:
- Septal leaflet
- Lateral leaflet
Right Atrium
19 Right Coronary
with calcification
Thebesian Valve
Heart
1000 µm

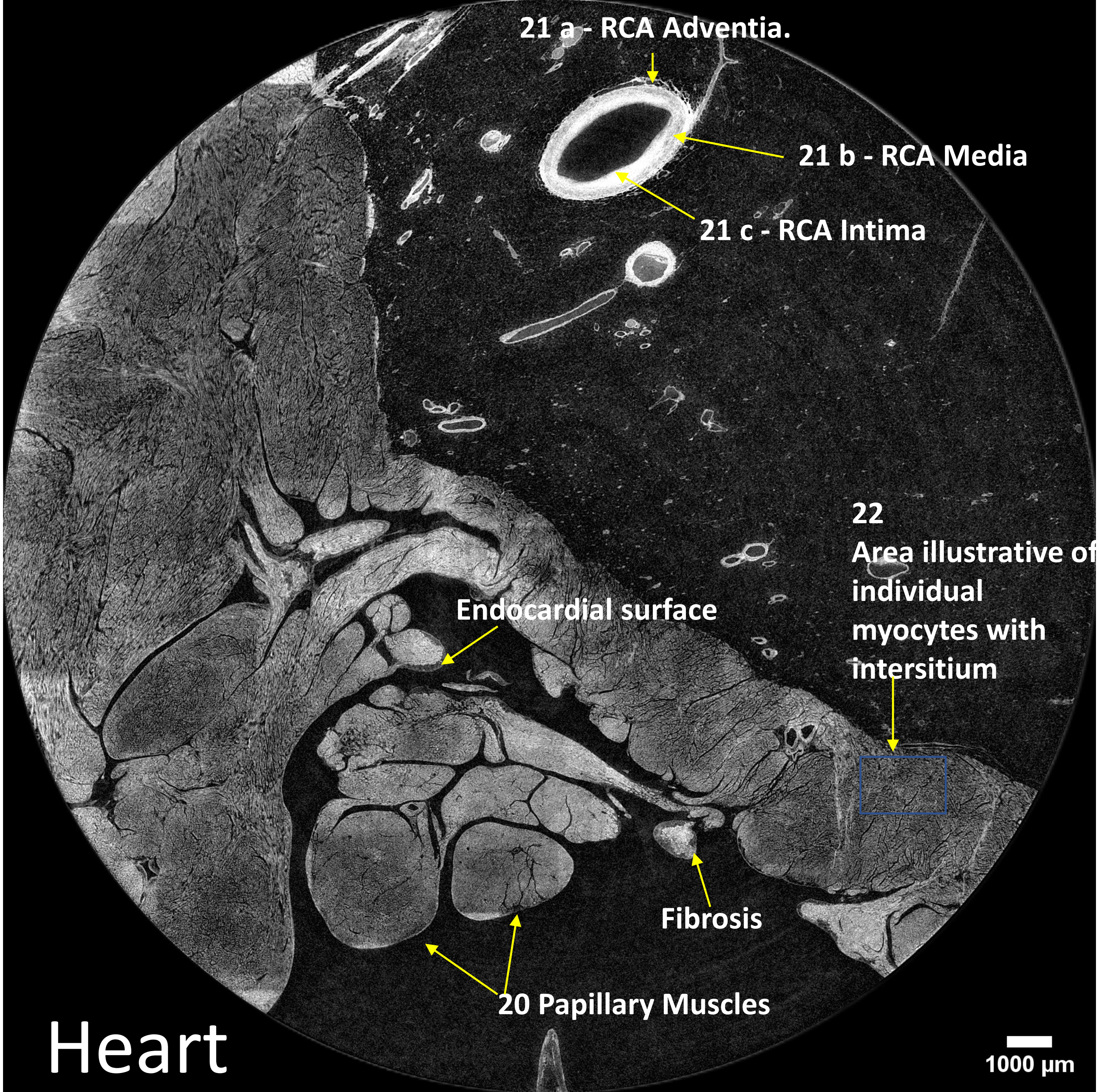
21 a - RCA Adventia.
21 b - RCA Media
21 c - RCA Intima
22
Area illustrative of
individual
myocytes with
intersitium
Endocardial surface
Fibrosis
20 Papillary Muscles
1000 µm

Heart

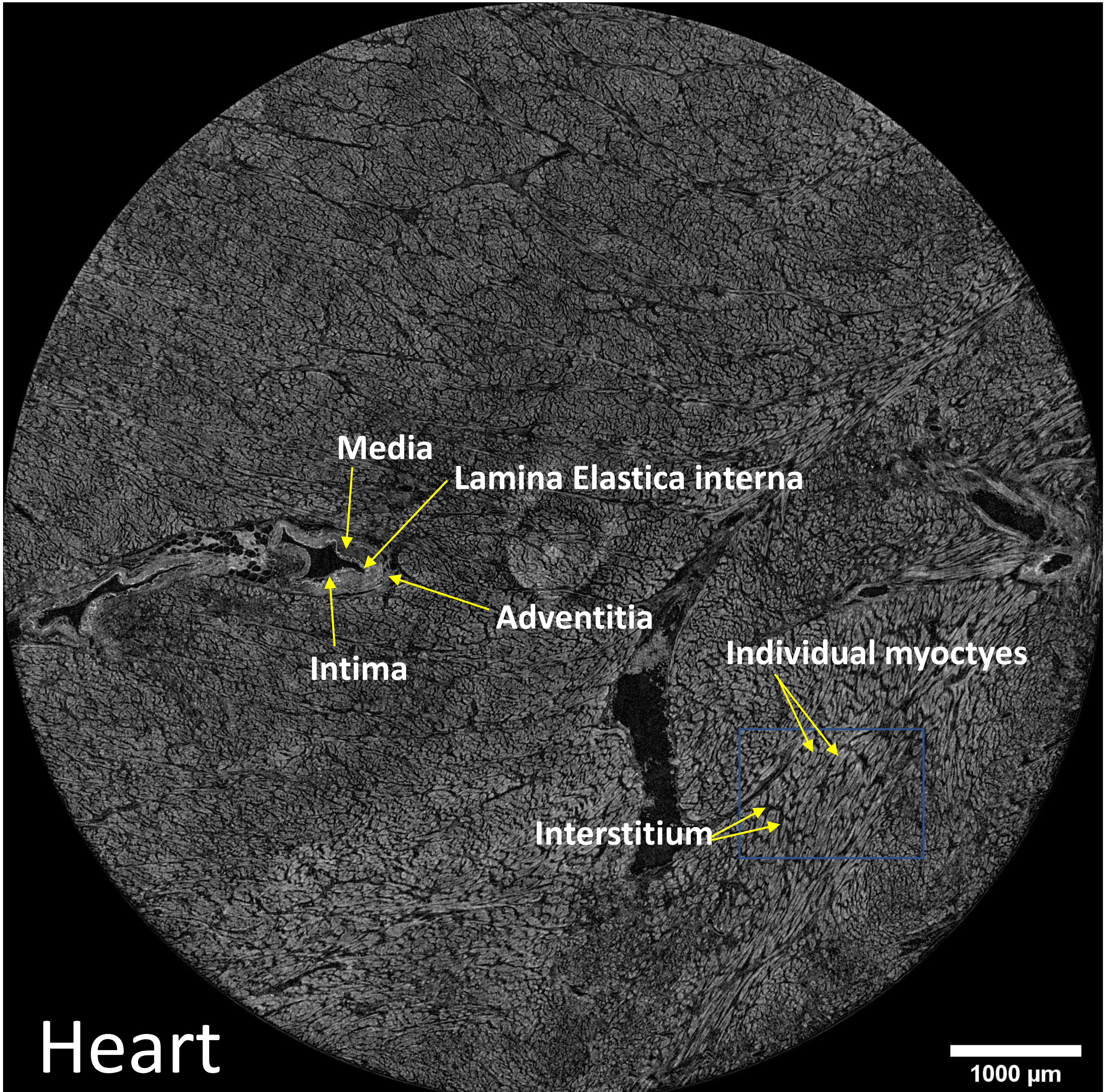
Media
Lamina Elastica interna
Adventitia
Intima
Individual myoctyes
Interstitium
Heart
1000 µm

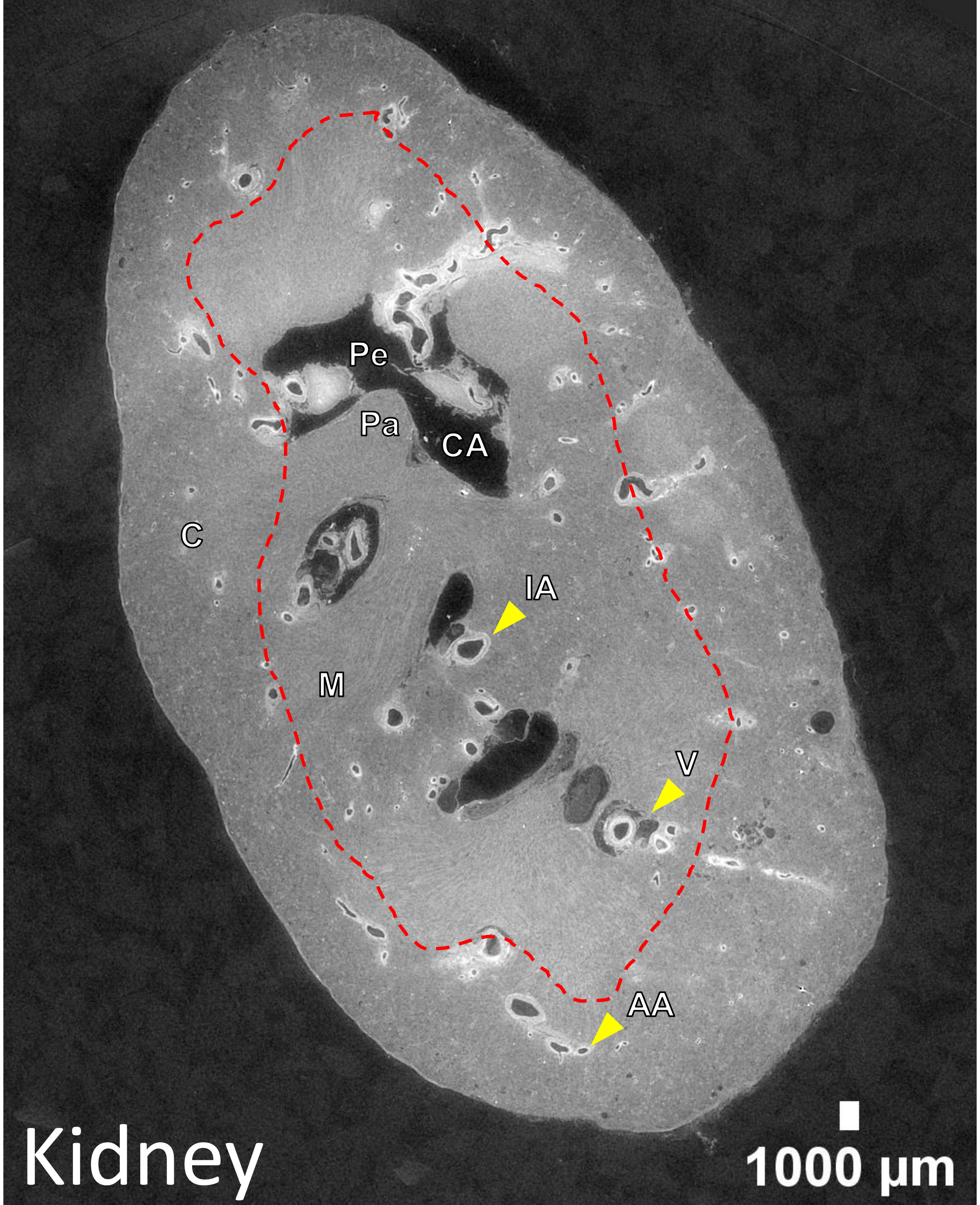


AA: Arcuate artery
C: Cortex
CA: Major calyx
IA: Interlobar artery
M: Medulla
Pa: Papilla
Pe: Pelvis
V: Venule
Dotted red line = corticomedullary junction

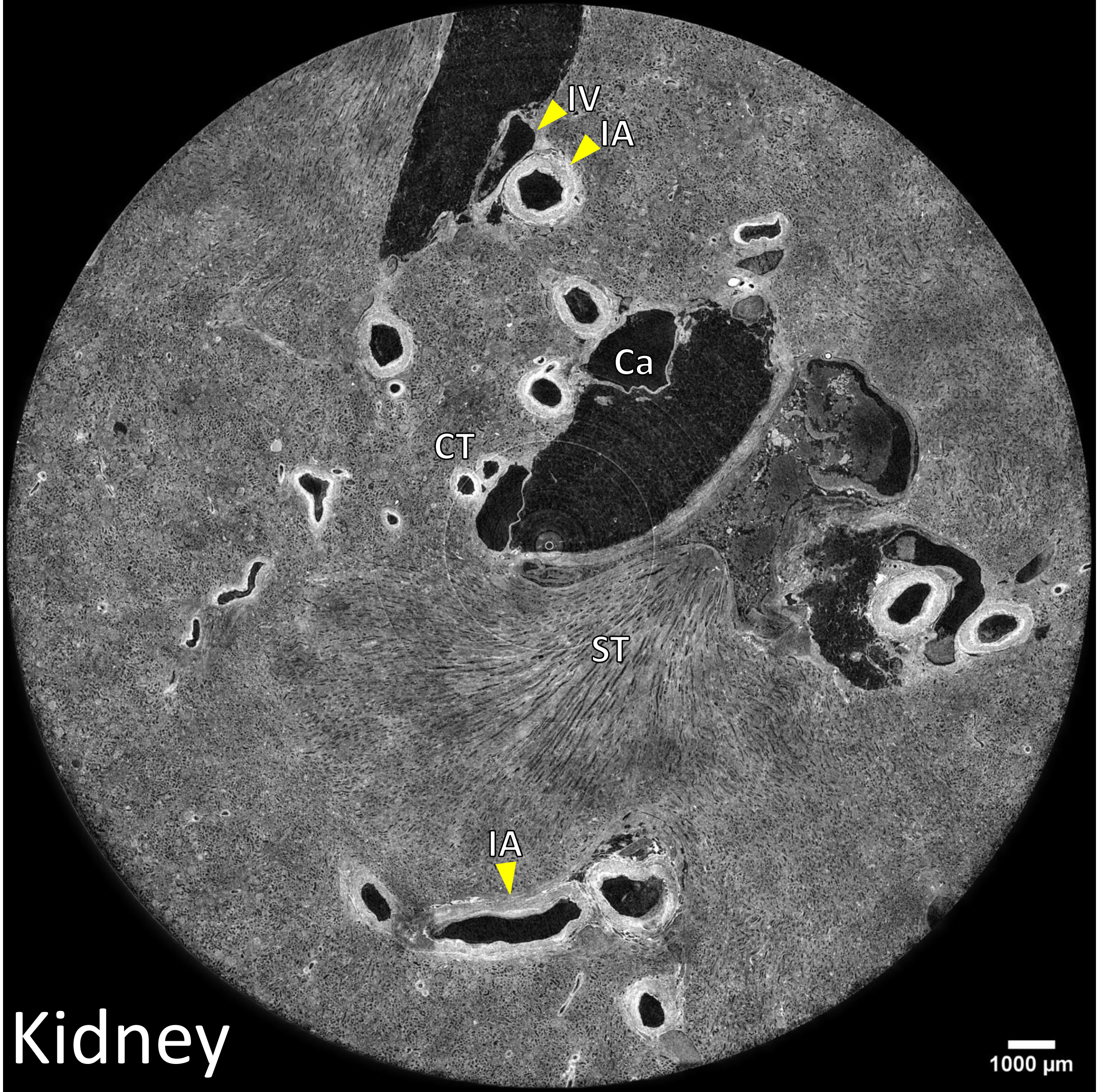


Ca: Minor calyx
CT: Convoluted tubules
IA: Interlobar artery
IV: Interlobar vein
ST: Straight tubules

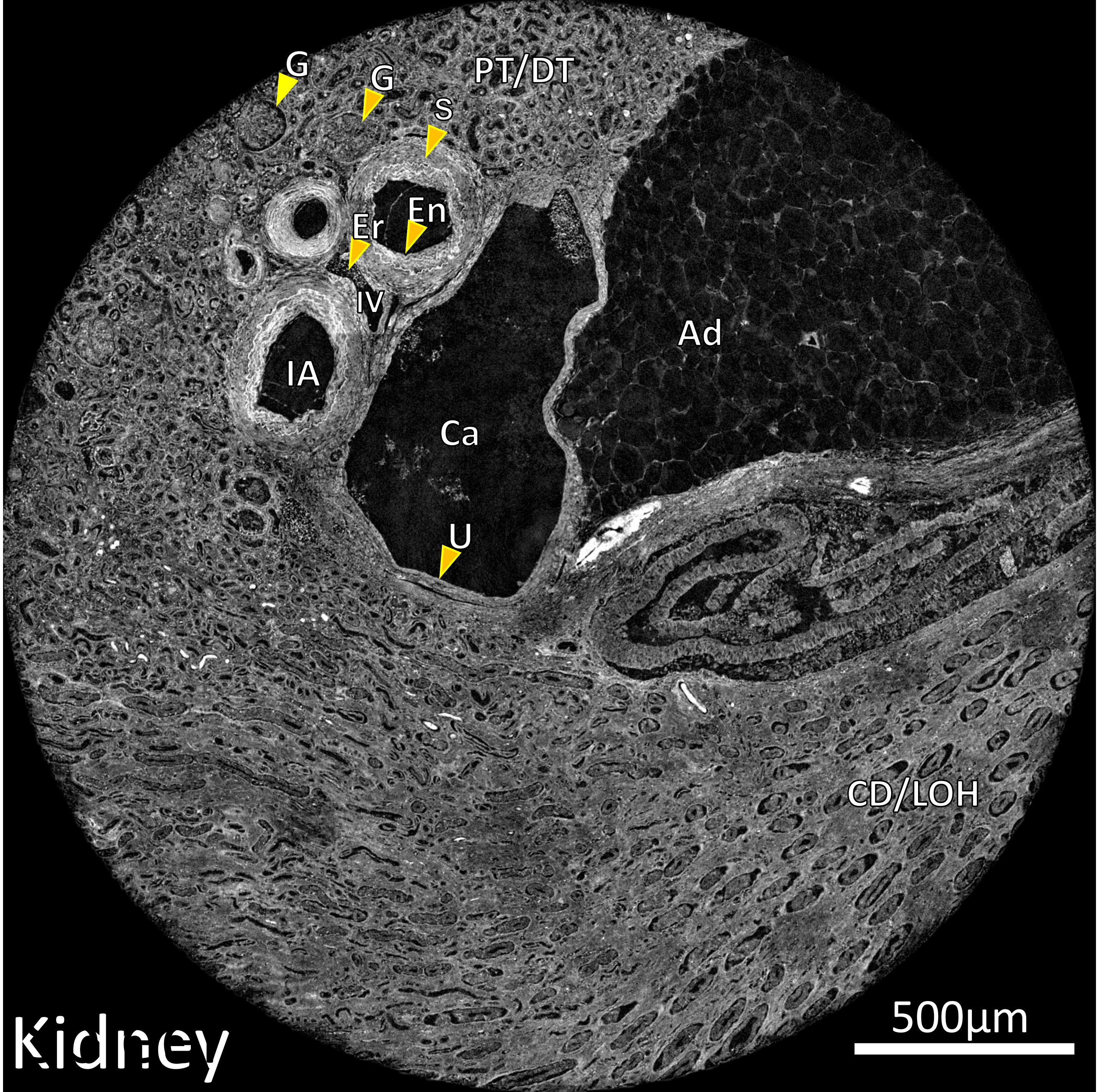


Ad: Adipocyte
Ca: Minor calyx
CD/LOH: Collecting duct / loop of Henle
En: Endothelium
Er: Erythrocyte
G: Glomerulus
IA: Interlobar artery
IV: Interlobar vein
PT/DT: Proximal tubule / distal tubule

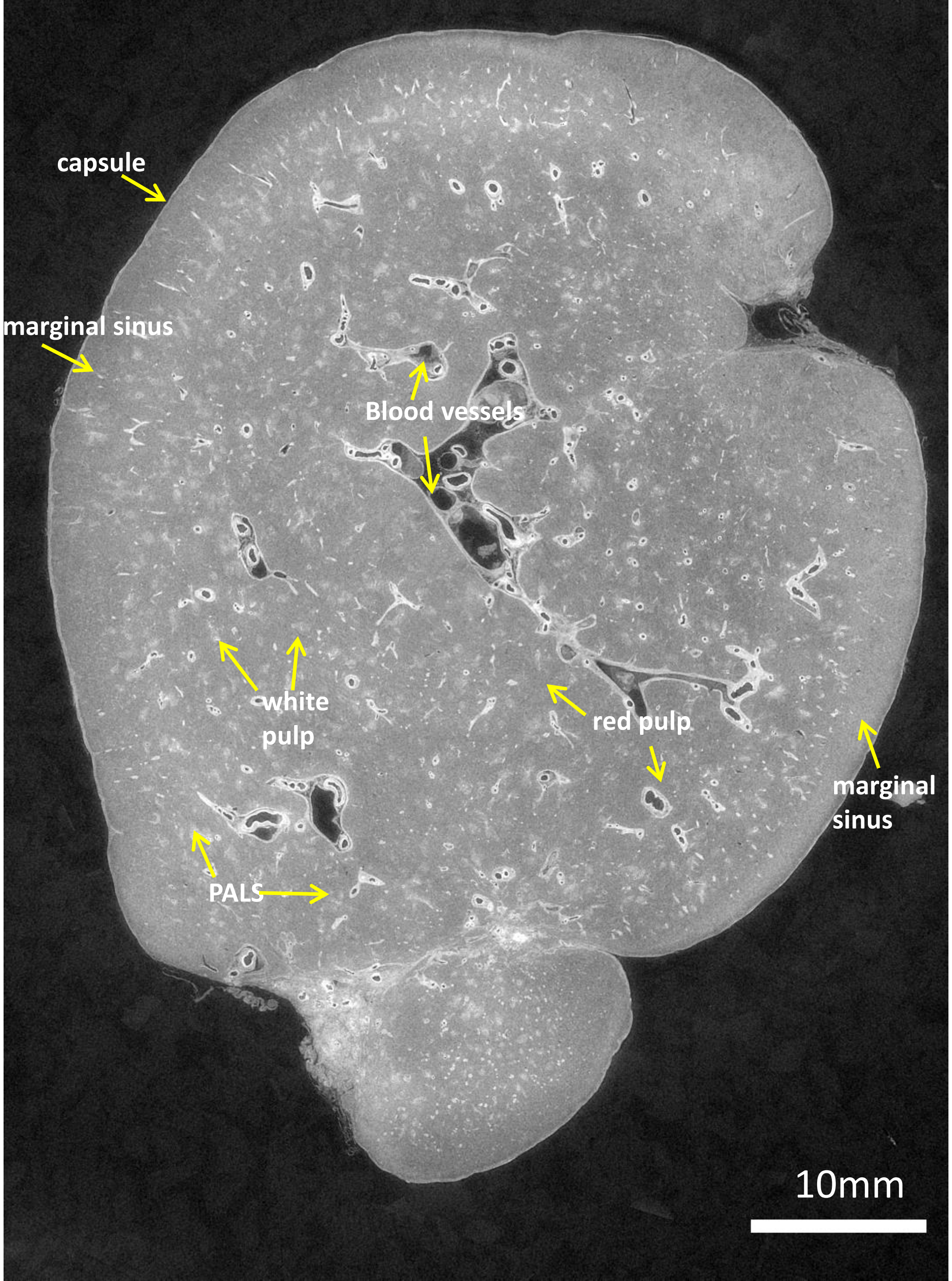
capsule
marginal sinus
Blood vessels
white pulp
red pulp
marginal sinus
PALS
10mm

### Spleen

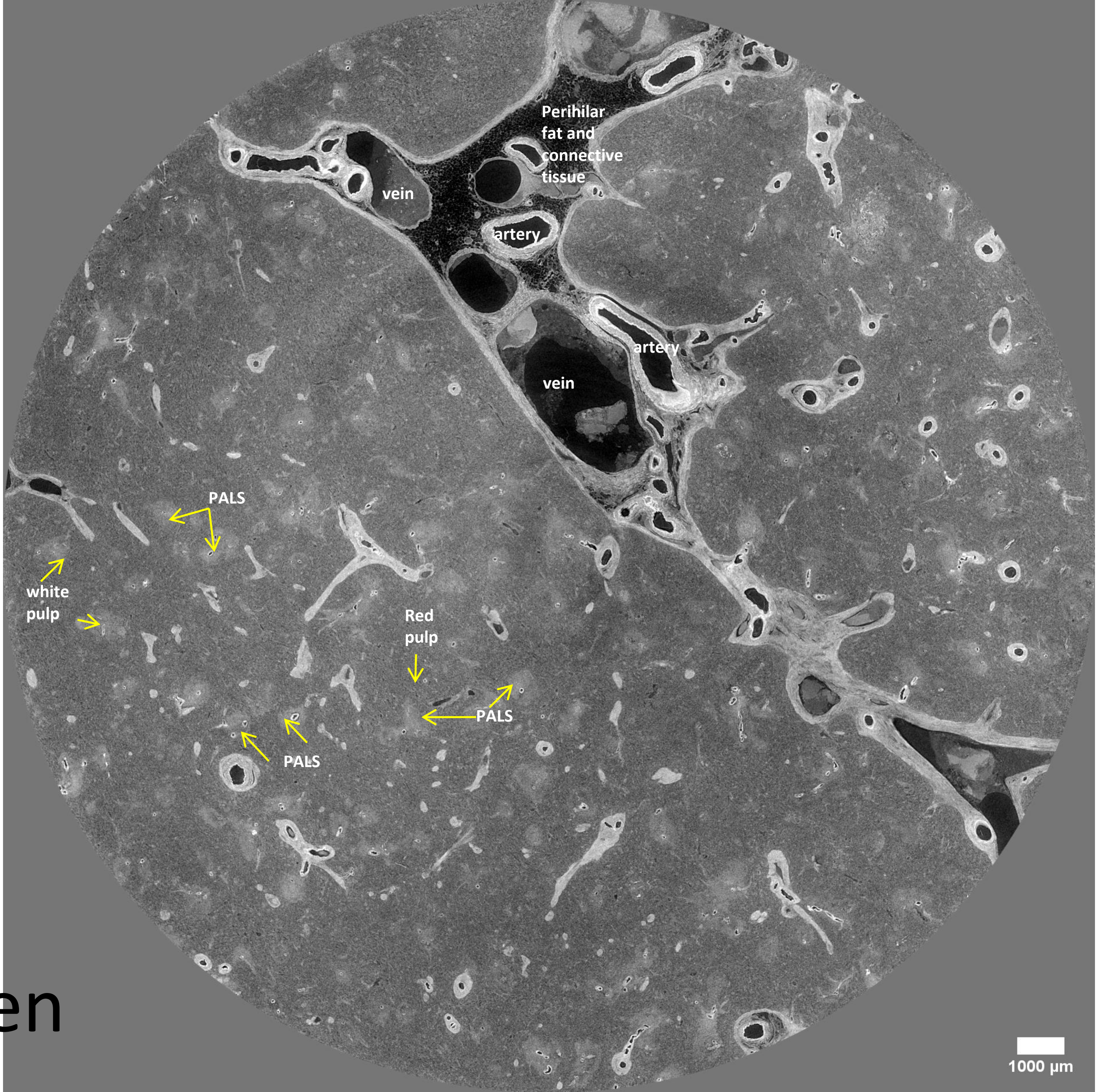
Perihilar fat and connective tissue
vein
artery
artery
vein
PALS
white pulp
Red pulp
PALS
PALS
1000 µm

### Spleen

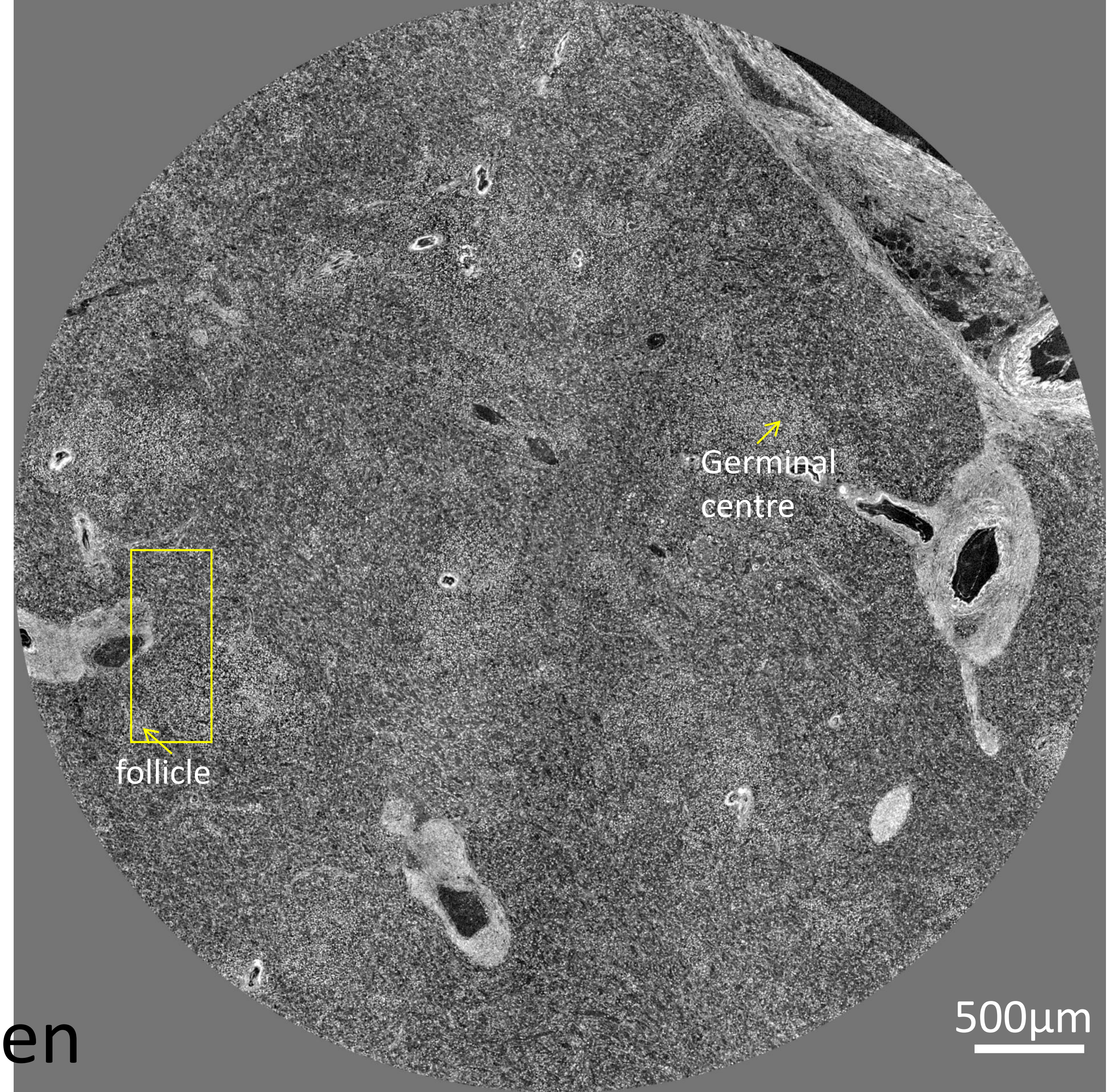


### Spleen
