## Supplementary information for "Multiscale three-dimensional imaging of intact human organs down to the cellular scale using hierarchical phase-contrast tomography"

### 1. Beam configuration

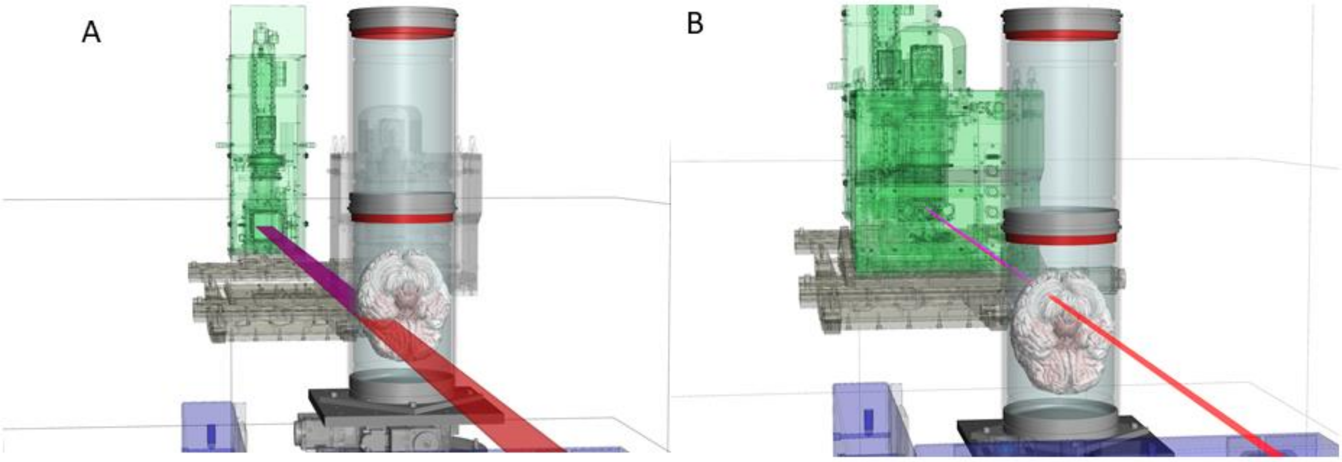


**Supplementary Figure 1:** Beam line configuration of human brain sample. Two containers (upper reference container and lower container holding the human brain sample) and the beam (red/magenta) are displayed in both images. A) 25 µm per voxel whole organ scan using the dzoom optic (green structure). B) a 2.5µm or lower resolution scan using the zoom optic (green structure).

### 2. Region selections

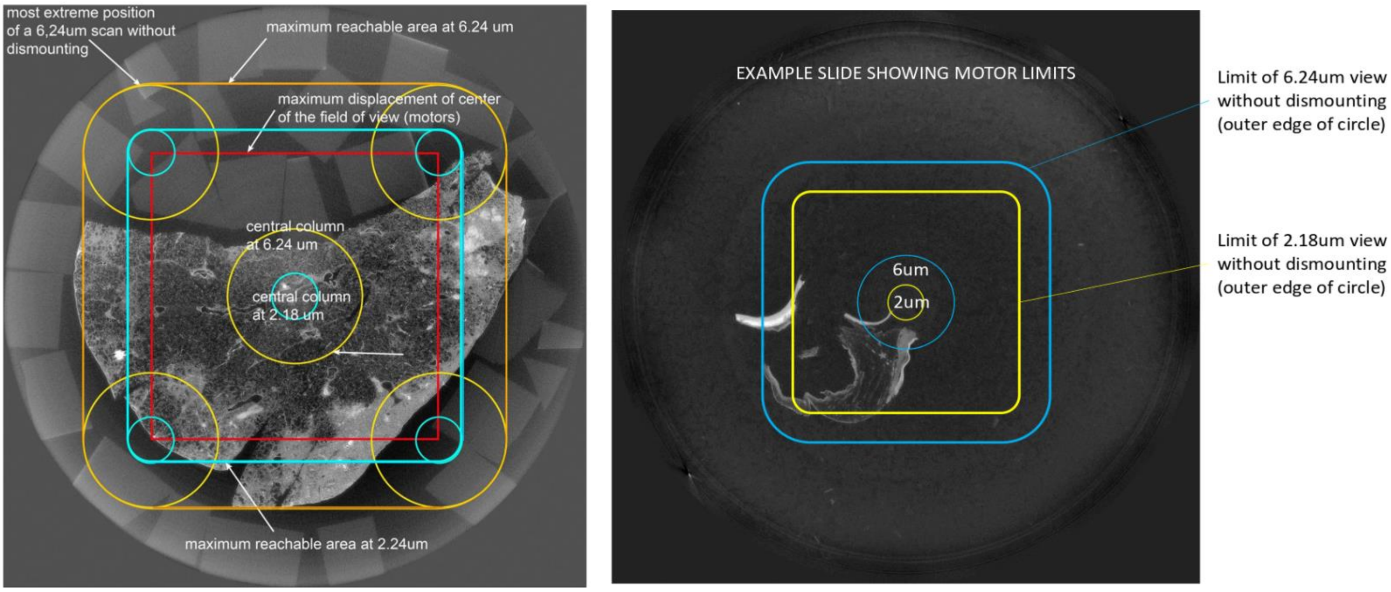


**Supplementary Figure 2:** Motor limits and VOI selection in the half (left) and quarter (right) acquisition modes.

**3 X-ray dose calculations**

The X-ray dose delivered to the surface of the sample with HiP-CT is much higher than for biopsied tissue and is estimated (as water equivalent surface dose) through a series of measurements with a dosimeter that have been used to develop a dose rate estimator for any beam configuration used on BM05, as shown in (**Supplementary Data 1**). The dose integrated on a small volume of interest in local tomography is lower than the total delivered dose, as a large part of the incoming X-rays is absorbed in the surrounding tissues and mounting media. Nevertheless, for comparable signal level, the dose will always be higher in HiP-CT than when scanning core biopsies, because the signal has to cross a lot of material before reaching the detector. The necessary total delivered dose to the sample to reach similar level of quality in HiP-CT compared to core biopsies is then about 4 times higher, for integrated dose in the VOI of typically twice higher. The main consideration for dose with ex vivo samples is to prevent bubble formation, which causes artefact and prevents ridged registration. These bubbles appear when a given level of integrated dose is reached. Once started, each new scan will increase the problem, even if done in a different location. the only way to recover the sample and make it suitable for new scans is to perform a new vacuum degassing. The tolerance of the sample to high level of dose is extremely dependant of the initial degassing level. During the early phase of this project, it appeared that careful degassing makes the sample able to handle total dose at least 10 times higher than without degassing. After a series of scans, putting the sample at 5 degrees for typically two days if no bubbling occurred helps a lot to prevent bubbling at the next scanning session.  Tissue damage may also be caused by excessive X-ray dose. At present, we have shown that even at the highest dosed areas (1.3µm columns that fall entirely within 6µm columns) histology with standard H&E can be performed and shows no obvious morphological damage (**Figure 3C**). Further evidence is shown in **Supplementary Figure 3** where the H&E stained large kidney section is shown with the outline of the scanned regions at 1.3µm and 6µm overlayed in green and blue respectively, there is no visible change in tissue morphology across these borders. Furthermore, immunohistochemistry (IHC) was performed on a Control and COVID lung biopsy sample (**Supplementary Figure 4**) indicating that neither the tissue pre-processing nor the X-ray dose (lower for biopsy samples) appear to adversely affect the expected staining pattern for a panel of commonly used IHC markers. Whilst we have not yet performed IHC on highest dose areas (1.3 µm within a HiP-CT scanned organ, these data strongly suggest that HiP-CT is compatible with IHC). The only cases that led to visible tissue damages occurred during the first period of development of HiP-CT when the acquisition system stopped for several hours.

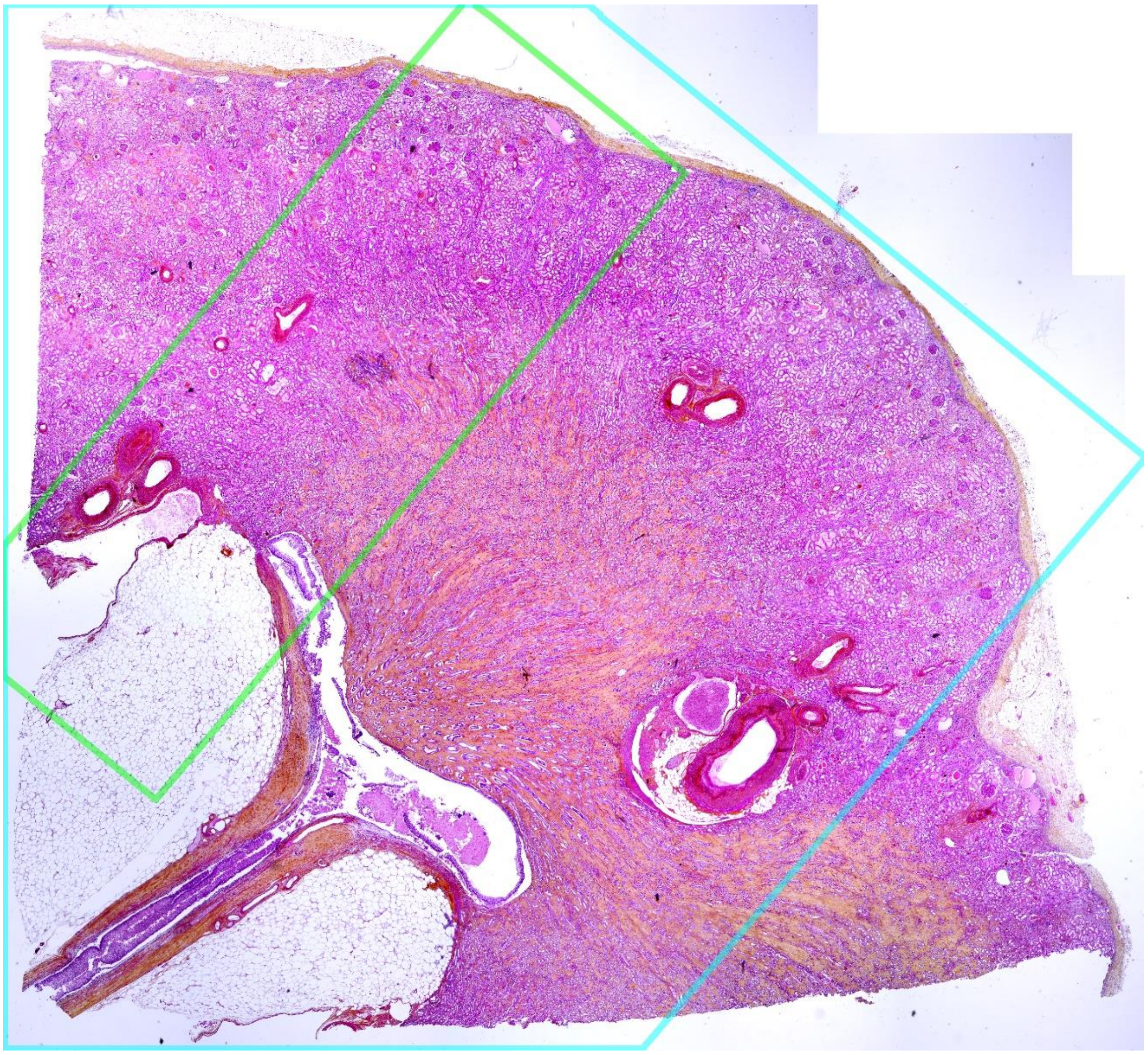

**Supplementary Figure 3.** Large scale view of Haemotoxylin and Eosin (H&E) stained slice of the kidney with outlines of the 6µm (blue) and 1.3µm (green) scanning regions shown. There is no visible differences between these regions.

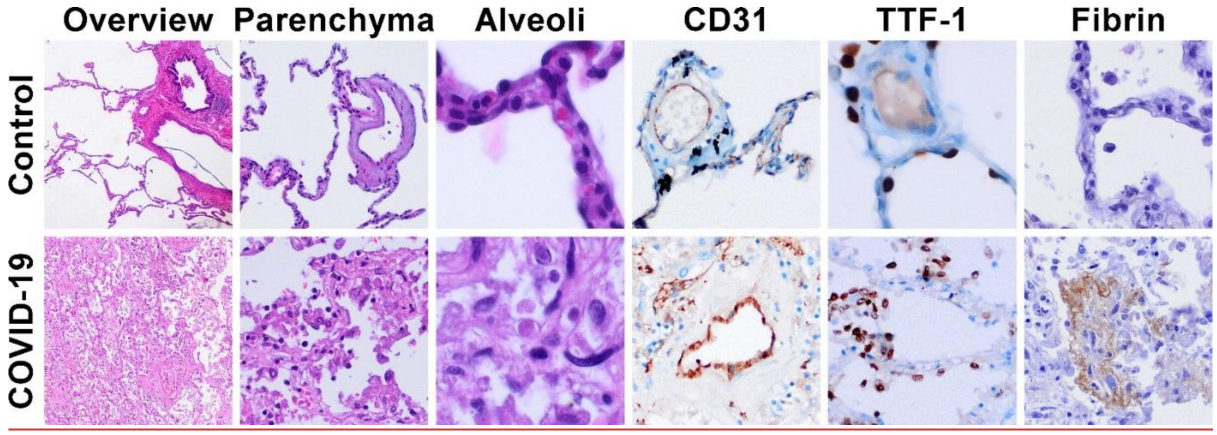


**Supplementary Figure 4:** H&E and IHC staining of COVID-19 lung biopsy and control lung biopsy after HiP-CT imaging. H&E staining of overview, parenchyma and alveoli in columns 1-3; CD31 (column 4), Thyroid transcription factor 1 (TTF-1 (column 5)) and Fibrin staining (column 6).

**Quantitative image Analysis: Machine Learning and Radiomics**

At the 6µm and 25µm scales we have begun to apply two techniques that have been transformational to medical image analysis in recent years: machine-learning and radiomics [1]. Supervised machine-learning techniques (ML), particularly convolutional neural networks (CNNs) can be trained to perform specific image processing tasks with as high an accuracy as human experts [2]. A CNN requires labelled datasets from which to iteratively train to make an image or pixel-wise 'decision' based on intrinsic features of the data. The strength of ML for HiP-CT is that whilst training is computationally intensive, once trained, the CNN can be applied to large datasets with relative ease. Training a successful CNN is dependent on having large well labelled ground truth datasets. As HiP-CT is a new modality, training datasets do not currently exist, and the expertise to perform the labelling as well as a choice of label categories required development. Through collaboration between image analysts, synchrotron scientists, radiologists and pathologists we trialled a number of labelling strategies and label categories to reach consensus between features that could be reliably manually distinguished and were of clinical interest. Our final strategy utilised the 6µm imaging datasets (**Supplementary Figure 5Ai**) and classified voxels into one of four categories: vessels, free air space, obstructed airspaces and unclassified tissue (mainly highly consolidated areas where no visible structures are discernible)  The 6µm dataset facilitated segmentation as it allows simultaneously viewing of the same area of lung at both greater detail (2µm) and with full organ context (25µm), which was important for distinguishing small, obstructed airways and vessels, particularly in damaged regions. To create training and validation, datasets of 5 VOIs (162x152x880 voxels) were chosen to reflect the range of tissue structures and damage seen throughout the COVID-19 lung (**Figure 4Ciii**). We then trained a 3D CNN developed for application on large 3D datasets (https://github.com/natalie11/tUbeNet). Training was run for 500 epochs (~48hrs); **Figure 5Cii** show training progress (loss and accuracy for over time). Both the accuracy and the loss have clearly plateaued by 500 epochs and whilst the curves for the validation dataset broadly follow a similar trend, there is a large separation between the training and validation curves, indicating that the training dataset must be enlarged further.

Radiomics is the extraction of quantitative image data or features including first order statistics: i.e. those based on the histogram of the image intensity values and textural descriptors - local patterns of intensity values. Radiomics has been widely applied to CT and PET medical imaging as well as to MRI [3].  There are many ways in which image texture can be represented mathematically and hence there are many radiomics features reflecting this. The features are grouped by the matrices on which they are calculated: Grey level co-occurance matrix (GLCM), Grey level run matrix (GLRM), Grey level size zone matrix (GLSZM) and Grey level dependence matrix (GLDM), all representing mathematical descriptions of which can be found in the Radiomics literature [3].  The output from a Radiomics analysis is a large panel of image features the user has chosen to include. These features amount to a reduction of the image parameter space and hence are amenable to parametric analyses such as principle component analysis or other clustering approaches. In recent years radiomics analyses have been used to combine imaging data with other types of clinical or genomic data to aid in diagnosis particularly in relation to lung cancer [3].

We chose to apply radiomics analysis to the 25µm data using the secondary pulmonary lobule as the structural unit. Secondary pulmonary lobules are discrete structures of the lung separated by interlobular septae and contain acini. These lobules cannot usually be distinguished by clinical CT in human patients due to limitations in spatial resolution and there are no studies to the authors knowledge that map secondary pulmonary lobule structures across the whole human lung. A striking finding from the 25µm HiP-CT dataset in COVID-19 was the large variation in structural deterioration between adjacent pulmonary lobules. The two lobules shown in **Figure 4B** adjoin one another but have very different structural appearance. Lobule 1 (yellow) shows clear open air spaces and wide preservation of alveolar structure, while lobule 2 (blue) shows large areas of consolidation and widespread loss of alveolar structure. **Figure 4Diii** shows the fold change in radiomics feature values between the two lobules for 89 different features (listed below). Features with a fold change higher than two standard deviations (and not confounded by volume) are labelled. Skewness is positive in lobule 1 (yellow) indicating that the mass of the intensity distribution is shifted to lower pixel intensities; skewness becomes negative, i.e. shifted towards higher pixel intensities, in lobule 2 (blue). Grey Level non-Uniformity is highlighted in each of the different feature groups. Lower values of this feature indicate more uniform tissue grey values, seen in lobule 1 (yellow) and less uniform in lobule 2 (blue). Whilst some of these individual parameters may be of interest, performing radiomics analysis on many more lobules will enable automated clustering analyses and correlation radiological terms e.g. consolidation to radiomic features, and potentially in the future correlation to clinical CT.

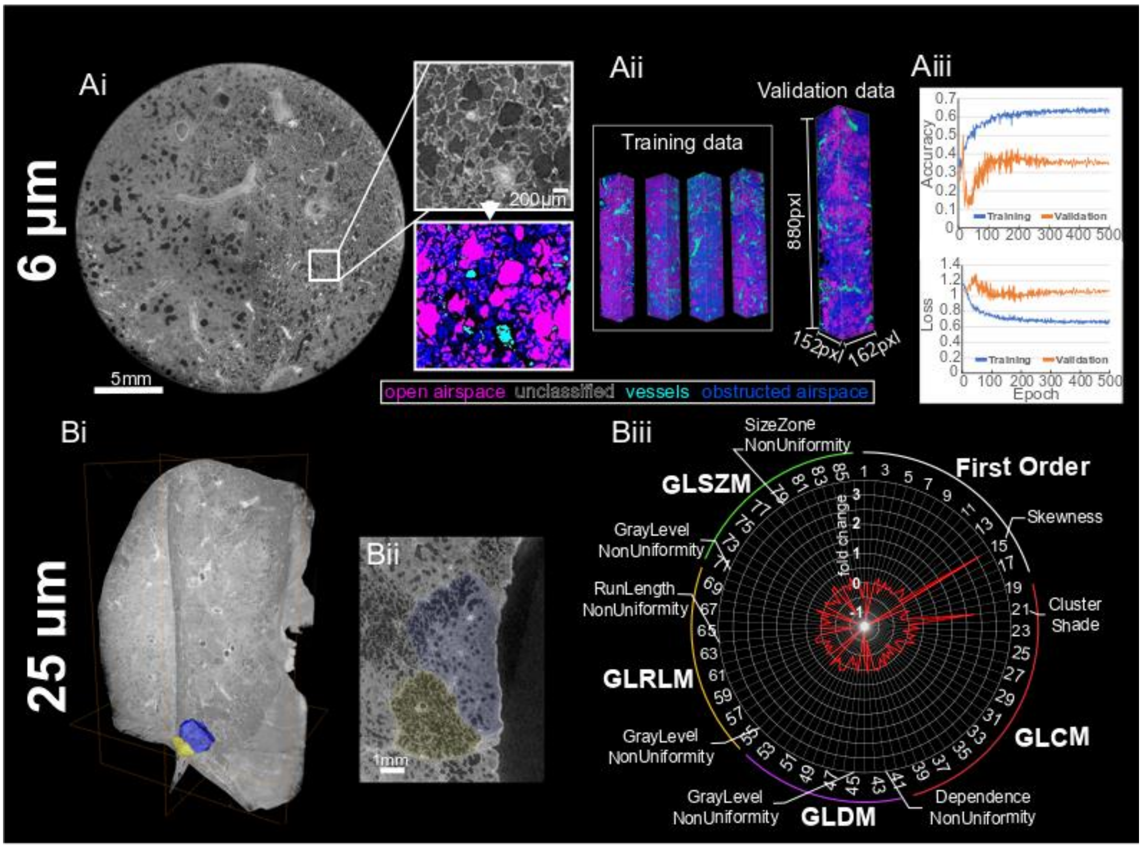


*Ai) Shows a typical slice from 6 µm COVID-19 dataset with a white box denoting a small volume which has been manually segmented. Free airspace (magenta), vessels (cyan), obstructed airspace (blue), and unclassified tissue (black) were trained on subvolumes. Aii) The training data and validation data were compiled from the 5 subvolumes (162x157x880pxls )which were chose to reflect the range of lung damage and structures within the whole dataset. Aiii) Loss and accuracy of the network for the training and validation datasets with increasing epochs. B) Analysis of COVID lung lobe at 25µm voxels using radiomics analysis based on the secondary pulmonary lobule. Bi) shows orthogonal slices through the lung lobe with two adjoining secondary pulmonary lobules segmented in blue and yellow. Bii) 2D section through the lung lobe with two secondary lobules highlighted in coloured overlays. The yellow lobule is less severely affected than the blue one. Biii) Radar plot*

*showing the outcome of comparative radiomics for the two lobules. 86 radiomics parameters across were included and the fold change between the yellow and blue lobule for each parameter calculated (yellow-blue)/yellow. Highlighted parameters include skewness (positive in the yellow lobe but becomes negative i.e. shifter towards higher pixel intensities in the more damaged lobe. Grey Level non-Uniformity is highlighted in several feature groups. Lower values of this feature indicate more uniform tissue grey values, indicating that the more severely affected lobule is less uniform.*

1. *original_firstorder_10Percentile*
2. *original_firstorder_90Percentile*
3. *original_firstorder_Energy*
4. *original_firstorder_Entropy*
5. *original_firstorder_InterquartileRange*
6. *original_firstorder_Kurtosis*
7. *original_firstorder_Maximum*
8. *original_firstorder_MeanAbsoluteDeviation*
9. *original_firstorder_Mean*
10. *original_firstorder_Median*
11. *original_firstorder_Minimum*
12. *original_firstorder_Range*
13. *original_firstorder_RobustMeanAbsoluteDeviation*
14. *original_firstorder_RootMeanSquared*
15. *original_firstorder_Skewness*
16. *original_firstorder_TotalEnergy*
17. *original_firstorder_Uniformity*
18. *original_firstorder_Variance*
19. *original_glcm_Autocorrelation*
20. *original_glcm_JointAverage*
21. *original_glcm_ClusterProminence*
22. *original_glcm_ClusterShade*
23. *original_glcm_ClusterTendency*
24. *original_glcm_Contrast*
25. *original_glcm_Correlation*
26. *original_glcm_DifferenceAverage*
27. *original_glcm_DifferenceEntropy*
28. *original_glcm_DifferenceVariance*
29. *original_glcm_JointEnergy*
30. *original_glcm_JointEntropy*
31. *original_glcm_Imc1*
32. *original_glcm_Imc2*
33. *original_glcm_Idm*
34. *original_glcm_Idmn*
35. *original_glcm_Id*
36. *original_glcm_Idn*
37. *original_glcm_InverseVariance*
38. *original_glcm_MaximumProbability*
39. *original_glcm_SumEntropy*
40. *original_glcm_SumSquares*
41. *original_gldm_DependenceEntropy*
42. *original_gldm_DependenceNonUniformity*
43. *original_gldm_DependenceNonUniformityNormalized*
44. *original_gldm_DependenceVariance*
45. *original_gldm_GrayLevelNonUniformity*
46. *original_gldm_GrayLevelVariance*
47. *original_gldm_HighGrayLevelEmphasis*
48. *original_gldm_LargeDependenceEmphasis*
49. *original_gldm_LargeDependenceHighGrayLevelEmphasis*
50. *original_gldm_LargeDependenceLowGrayLevelEmphasis*
51. *original_gldm_LowGrayLevelEmphasis*
52. *original_gldm_SmallDependenceEmphasis*
53. *original_gldm_SmallDependenceHighGrayLevelEmphasis*
54. *original_gldm_SmallDependenceLowGrayLevelEmphasis*
55. *original_glrlm_GrayLevelNonUniformity*
56. *original_glrlm_GrayLevelNonUniformityNormalized*
57. *original_glrlm_GrayLevelVariance*
58. *original_glrlm_HighGrayLevelRunEmphasis*
59. *original_glrlm_LongRunEmphasis*
60. *original_glrlm_LongRunHighGrayLevelEmphasis*
61. *original_glrlm_LongRunLowGrayLevelEmphasis*
62. *original_glrlm_LowGrayLevelRunEmphasis*
63. *original_glrlm_RunEntropy*
64. *original_glrlm_RunLengthNonUniformity*
65. *original_glrlm_RunLengthNonUniformityNormalized*
66. *original_glrlm_RunPercentage*
67. *original_glrlm_RunVariance*
68. *original_glrlm_ShortRunEmphasis*
69. *original_glrlm_ShortRunHighGrayLevelEmphasis*
70. *original_glrlm_ShortRunLowGrayLevelEmphasis*
71. *original_glszm_GrayLevelNonUniformity*
72. *original_glszm_GrayLevelNonUniformityNormalized*
73. *original_glszm_GrayLevelVariance*
74. *original_glszm_HighGrayLevelZoneEmphasis*
75. *original_glszm_LargeAreaEmphasis*
76. *original_glszm_LargeAreaHighGrayLevelEmphasis*
77. *original_glszm_LargeAreaLowGrayLevelEmphasis*
78. *original_glszm_LowGrayLevelZoneEmphasis*
79. *original_glszm_SizeZoneNonUniformity*
80. *original_glszm_SizeZoneNonUniformityNormalized*
81. *original_glszm_SmallAreaEmphasis*
82. *original_glszm_SmallAreaHighGrayLevelEmphasis*
83. *original_glszm_SmallAreaLowGrayLevelEmphasis*
84. *original_glszm_ZoneEntropy*
85. *original_glszm_ZonePercentage*
86. *original_glszm_ZoneVariance*
